# Selective MAP1LC3C (LC3C) autophagy requires noncanonical initiation regulators and the paralog-specific C-terminal peptide

**DOI:** 10.1101/2021.02.19.431861

**Authors:** Megan E. Bischoff, Yuanwei Zang, Johnson Chou, Adam D. Price, Birgit Ehmer, Nicholas J. Talbot, Michael J. Newbold, Anurag Paul, Jun-Lin Guan, David R. Plas, Jarek Meller, Maria F. Czyzyk-Krzeska

## Abstract

LC3s are canonical proteins necessary for the formation of autophagosomes. We have previously established that two paralogs, LC3B and LC3C, have opposite activities in renal cancer, with LC3B playing oncogenic role and LC3C tumor suppressing role. LC3C is an evolutionary late gene, present only in higher primates and humans. Its most distinct feature is a C-terminal 20 amino acid peptide cleaved in the process of glycine 126 lipidation. Here we investigated mechanisms of LC3C selective autophagy. LC3C autophagy requires noncanonical upstream regulatory complexes that include ULK3, UVRAG, RUBCN, PIK3C2A, and a member of ESCRT, TSG101. We established that Postdivision Midbody Rings (PDMBs) implicated in cancer stem cell regulation are direct targets of LC3C autophagy. LC3C C-terminal peptide is necessary and sufficient to mediate LC3C-dependent selective degradation of PDMBs. This work establishes a new noncanonical human-specific selective autophagic program relevant to cancer stem cells.

## Introduction

Macroautophagy (autophagy), an essential pathway leading to lysosomal degradation of intracellular organelles, long-lived proteins, and RNAs, is a tightly regulated cellular process of major importance in cancer (White, 2015). Oncogenic autophagy provides nutrients, particularly when the supply from extracellular microenvironments is limited, and contributes to therapy resistance. The tumor suppressing activity of autophagy compensates for pathologic alterations to organelles/proteins, eg. aged mitochondria generating reactive oxygen species and the resulting DNA damage. Thus, biological activity of autophagy requires selectivity in the recognition of a specific cargo to be targeted for degradation under particular conditions. Selectivity in autophagy results from the activity of cargo receptor proteins that are able to recognize cargo in ubiquitin-dependent or independent manner and to bind to the ATG8 family of small proteins that are inserted into the autophagosome membrane (Galluzzi et al., 2017; Zaffagnini and Martens, 2016). Mammalian homologues of ATG8 include three paralogs of Microtubule Associated Protein 1 Light Chain 3 Alpha, Beta, and Gamma (LC3A, LC3B, and LC3C) and members GABA Type A Receptor-Associated Proteins (GABARAPs). The ubiquitylated cargo binds to the ubiquitin-binding domains (UBA) present on several canonical cargo receptors (eg. p62, NBR1, CALCOCO2-3). In turn, these receptors bind to LC3s or GABARAPs through LC3 Interacting Regions (LIR) motifs. While there are examples of ubiquitin-independent selective autophagy, the mechanisms of cargo recognition are not well understood (Zaffagnini and Martens, 2016).

We established that LC3B- and LC3C-dependent autophagic programs have opposite functions in clear cell renal cell carcinoma (ccRCC), regulated by VHL, the main tumor suppressor lost in ccRCC (Hall et al., 2014; Mikhaylova et al., 2012). LC3B-mediated autophagy plays a key role in the progression of ccRCC and is inhibited by VHL (Hall et al., 2014; Mikhaylova et al., 2012). In contrast, LC3C autophagy has tumor suppressing activity in a renal cancer xenograft model and is positively regulated by VHL due to the inhibition of HIFs, which transcriptionally repress LC3C (Mikhaylova et al., 2012). Thus, VHL orchestrates a shift between tumor suppressing and oncogenic autophagic programs. Tumor suppressing activity of LC3C autophagy in renal cancer is further supported by the fact that LC3C activity is induced by folliculin (FLCN), another renal tumor suppressor (Bastola et al., 2013) and that it regulates of Met-RTK and HGF-stimulated migration and invasion (Bell et al., 2019). Other known LC3C functions, include xenophagy (von Muhlinen et al., 2012), piecemeal mitophagy (Le Guerroue et al., 2017), viral trafficking (Madjo et al., 2016), and exit of proteins from the endoplasmic reticulum (Stadel et al., 2015).

Midbody is a structure built at the intercellular bridge during cytokinesis, forming a platform for the complex process of abscission (White and Glotzer, 2012). After division, the postdivision midbody (PDMB) is inherited by one of the daughter cells in the case of asymmetric division, or by none, if the abscission happens on both sides of the midbody, and it is released into the extracellular space (Crowell et al., 2014; Fazeli and Wehman, 2017; Kuo et al., 2011). Asymmetrically inherited PDMB can undergo macroautophagy (Kuo et al., 2011; Pohl and Jentsch, 2009). PDMBs released outside the cell can be retained on the cell surface and then become phagocytosed (Crowell et al., 2014). In Drosophila, asymmetric inheritance of PDMB by female but not male germline stem cells is considered a platform for asymmetric segregation of molecules (Salzmann et al., 2014). In cancers, a high number of PDMBs promotes reprogramming towards pluripotency and increased tumorigenic potential in vitro (Kuo et al., 2011).

Here we investigated mechanisms contributing to the selectivity of the LC3C tumor suppressing autophagy. We determined that LC3C autophagic program requires noncanonical members in the upstream regulatory complexes including little known ULK3 as a part of FP200/ATG13/ULK complex, and UVRAG, RUBCN, and PIK3C2A as part of PI3K complex, as well as the cargo receptor, CALCOCO2. We identified PDMBs as specific LC3C cargo degraded by the lysosome, an implication that LC3C prevents reprogramming towards pluripotency. Most importantly, we show that LC3C C-terminal peptide is necessary and sufficient for the selectivity of LC3C-dependent degradation of PDMBs and for the assembly of the upstream noncanonical complexes with LC3C. We determined that the ESCRT-I member, TSG101, is required for the assembly of the upstream regulators with LC3C in a C-terminal peptide-dependent manner. The data establish a new mechanism of human specific, selective autophagy in the degradation of oncogenic cargo.

## Results

### LC3C and LC3B are independent autophagic programs

LC3C differs from LC3B/A in several important ways (Figure 1A). There is a 61% amino acid difference in the two N-terminal α-helices forming the LIR-binding domain as well as CLIR-binding motif that interacts with LC3C Interacting Region (CLIR) identified on CALCOCO2 cargo receptor (von Muhlinen et al., 2012). This amino acid difference is sufficient for generation of LC3C-specific antibodies (Figure S1A). LC3C has a 20 amino acid C-terminal peptide after G126 that undergoes lipidation during autophagosome formation. The peptide is highly conserved among primates (Figure S1B). LC3C, similarly to LC3B, accumulates in response to lysosomal inhibition using chloroquine (CQ) or bafilomycin A1 (Figures S1C and S1D), but LC3C and LC3B mainly form separate vesicles, with only a small fraction colocalized as shown in four different cell lines (Figure S1D). Importantly, serum starvation (0.1% serum in the media) causes fast, strong and long-lasting induction of LC3C mRNA accumulation, while this effect is significantly less strong in the case of LC3B (Figure S1E). The importance of LC3C in normal kidney epithelial cells is further underscored by the observation that immortalized human renal proximal tubule epithelial cells (RPTEC) show strong induction of LC3C, but not LC3B, in response to serum starvation (Figure S1F). Note that in the case of LC3C we detect only the LC3C form that accumulates in the presence of CQ or BAFA1 and represents the lipidated, LC3C-II form, while we do not detect full size LC3C or the LC3C-I. This likely indicates very fast conversion of the full size LC3C to the lipidated form and a mechanism of regulation that has other rate limiting steps than glycine lipidation. In addition to VHL, LC3C is also positively regulated by another renal cancer tumor suppressor, FLCN. Reexpression of FLCN in UOK 257 cells, which derive from a human Birt-Hogg-Dube patient and have lost endogenous FLCN, induces LC3C and represses LC3B flux, with only minor changes at the mRNAs levels (Figure S1G). Overall our data support different mechanisms of regulation and physiological activities of the LC3 paralogs.

**Figure 1.**
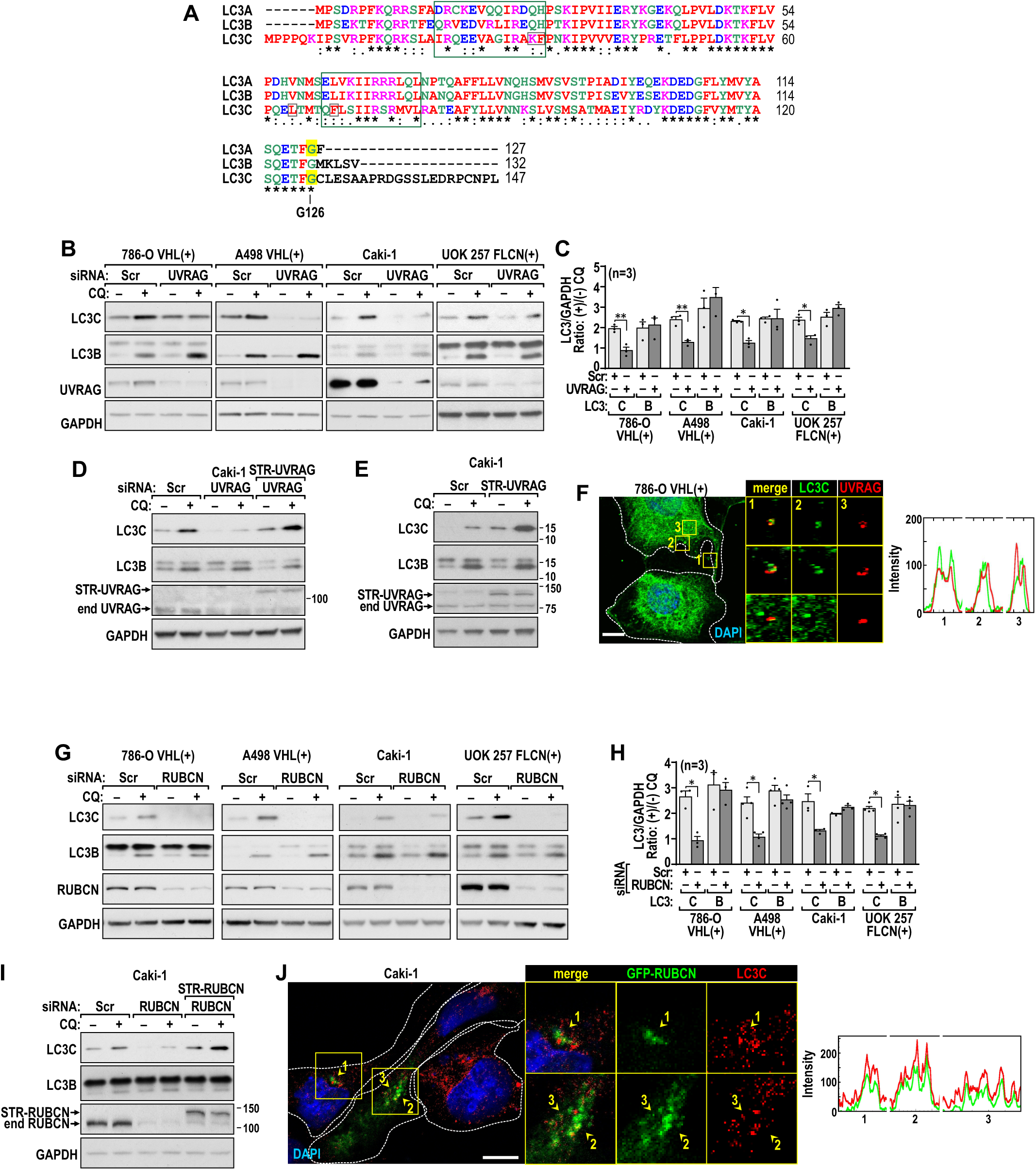
UVRAG and RUBCN regulate LC3C autophagy. (A) Sequence alignment of LC3 paralogs. At the bottom: (*)-conserved, (:)-conservative, (.)-non-conserved changes in amino acids composition. Large green boxes represent α-helices forming the LIR binding domain for the cargo receptors. Small gray boxes mark amino acids forming CLIR binding site; C-terminal G126 which undergoes lipidation is marked in yellow. (B) Immunoblots show decrease in the CQ dependent accumulation of LC3C in response to UVRAG knockdown in the indicated cell lines. (C) Quantification of the LC3C and B accumulation presented as the ratio of their protein levels normalized to GAPDH in CQ(+) vs. CQ(-) in cells expressing scrambled (Scr) or UVRAG siRNAs in the indicated cell lines. (D) Rescue of LC3C autophagic flux by reexpression of exogenous Strawberry tagged UVRAG (STR-UVRAG) to levels similar to that of endogenous UVRAG (end UVRAG). (E) Overexpression of STR-UVRAG induces LC3C autophagic flux. (F) STR-UVRAG colocalizes with endogenous LC3C. Split channels and RGB plots are shown below. Cells were treated with 100 µM CQ for 1 hr. (G) Immunoblots show decrease in the CQ dependent accumulation of LC3C in response to RUBCN knockdown in indicated cell lines. (H) Quantification of the LC3C and B accumulation presented as the ratio of their protein levels normalized to GAPDH in CQ(+) vs. CQ(-) in cells transfected with scrambled (Scr) or RUBCN siRNAs in the indicated cell lines. (I) Rescue of the LC3C autophagic flux by reexpression of exogenous Strawberry tagged RUBCN (STR-RUBCN) to levels similar to that of endogenous RUBCN (end RUBCN). (J) GFP-RUBCN colocalizes with endogenous LC3C. Split channels and RGB plots are shown. Cells were treated with 100 µM CQ for 1 hr. Means ± SEM are shown; P values from unpaired two tailed t-test. Scale bars: 10 µm.

### LC3C autophagy is regulated by noncanonical upstream complexes

In order to identify members of LC3C upstream regulatory complexes, we knocked down several members of ULK and PI3K-associated complexes and determined accumulation of LC3C when cells were grown in 0.1% serum for 48 hr and were not treated or treated with CQ for 1 hr before collection. We determined that BECN1 (Figure S1H) but not ATG14 (Figure S1I) was necessary for LC3C accumulation. ATG14-independent autophagy requires UVRAG in the PI3K-associated complex (Liang et al., 2006; Liang et al., 2008; Matsunaga et al., 2009). Indeed, knockdown of UVRAG inhibited LC3C autophagic flux, but was without significant effect on LC3B autophagy (Figures 1B and 1C). LC3C autophagic flux was rescued by reexpression of exogenous STR-UVRAG to the protein levels of endogenous UVRAG (Figure 1D) and was induced by overexpression of exogenous UVRAG (Figure 1E). Endogenous UVRAG colocalized with endogenous LC3C but not LC3B (Figures 1F, S1J and S1K). UVRAG had no effect on LC3C mRNA levels (Figure S1L).

UVRAG can function in autophagy and in a related process, LC3-Associated Phagocytosis (LAP) (Martinez et al., 2015). The activity of Rubicon (RUBCN) discriminates between these two UVRAG activities. RUBCN is an inhibitor of the BECN1/UVRAG complex in autophagy (Matsunaga et al., 2009), but an activator of the BECN1/UVRAG complex in LAP (Martinez et al., 2015). Knockdown of RUBCN inhibited LC3C autophagy (Figures 1G and 1H). Strawberry tagged-RUBCN (STR-RUBCN) rescued the effects of the knockdown (Figure 1I) and RUBCN colocalized with LC3C but not LC3B (Figure 1J). There were no effects of RUBCN knockdown on LC3C mRNA (Figure S1M).

PIK3C3 is required for canonical activation of autophagy. Surprisingly, LC3C autophagic flux was not affected by the knockdown of PIK3C3 (Figure S2A and S2B). Instead, we determined that a PI3K class II member, PIK3C2A, was necessary for LC3C autophagy (Figures 2A and 2B). The effect of PIK3C2A knockdown was rescued by reexpression of exogenous PIK3C2A (Figure 2C). Endogenous PIK3C2A colocalized with LC3C (Figure 2D) but not with LC3B (Figure S2C). Knockdown of PIK3C2A did not affect LC3C mRNA expression levels (Figure S2D). Further on we determined that LC3C colocalized with GFP-2xFyve, a reporter for PI3P, in a PIK3C2A- but not PIKC3C-dependent manner, supporting the role of this lipid and PIK3C2A in the formation of LC3C vesicles (Figures 2E, S2E and 2F). The formation of this complex was supported by co-immunoprecipitation of endogenous BECN1, RUBCN, PIK3C3A and LC3C with endogenous UVRAG in 786-O VHL(+) and Caki-1 cells (Figure 2G and 2H).

**Figure 2.**
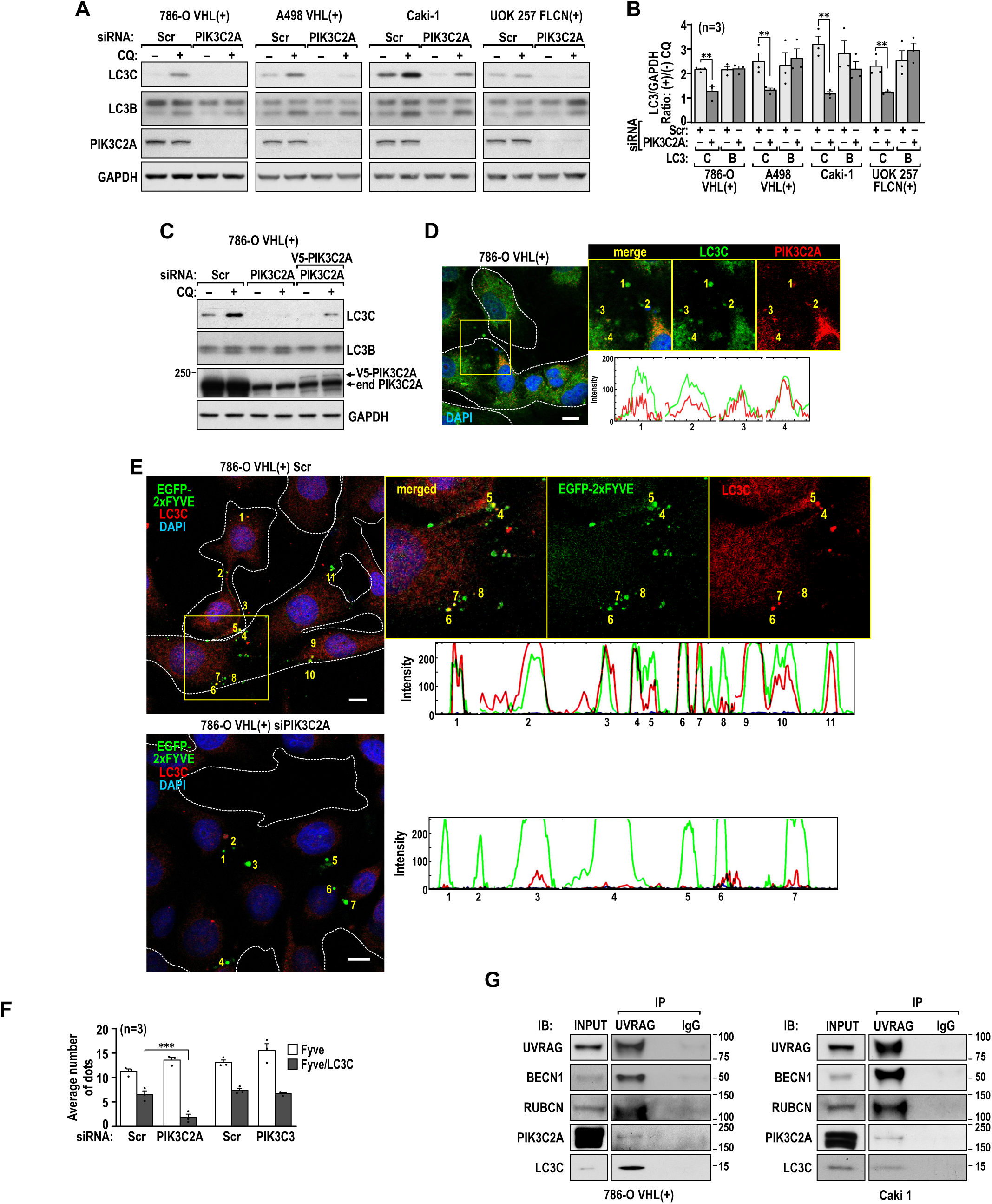
LC3C noncanonical upstream complexes include PIK3C2A rather than PIK3C3. (A) Immunoblots show decrease in the CQ dependent accumulation of LC3C in response to PIK3C2A knockdowns in the indicated cell lines. (B) Quantification of the LC3C and B accumulation presented as the ratio of their protein levels normalized to GAPDH in CQ(+) vs. CQ(-) in cells transfected with scrambled or PIK3C2A siRNAs in the indicated cell lines. (C) Rescue of the LC3C autophagic flux by reexpression of exogenous V5 tagged PIK3C2A. (D) PIK3C2A colocalizes with endogenous LC3C. Split channels RGB plots are shown. Cells were treated with 100 µM CQ for 1 hr. (E) Colocalization of LC3C with PI3P reporter, EGFP-2xFyve, in control cells and loss of this colocalization in cells with PIK3C2A knockdown. (F) Quantification of LC3C-EGFP-2xFyve colocalization in cells with PIK3C2A and PIK3C3 knockdowns. (G) Co-immunoprecipitation of endogenous BECN1, RUBCN, and PIK3C2A and LC3C with endogenous UVRAG in 786-O VHL(+) and Caki-1 cells. Means ± SEM are shown; P values from unpaired two tailed t-test. Scale bars: 10 µm.

Analysis of the role of FIP200/ATG13/ULK in LC3C lipidation, revealed that LC3C, similar to LC3B, required FIP200 (Figure S3A and S3B) and ATG13 (Figure S3C and S3D). In order to determine which ULK is involved in LC3C regulation, we measured expression levels of ULK1, ULK2 and ULK3 in all four cell lines (Figure S3E). Surprisingly, expression of ULK3 mRNA was by an order of magnitude higher than expression of ULK1 mRNA, while expression of ULK2 was found only in Caki-1 and UOK 257 FLCN(+), while it was extremely low in the other two cell lines (Figure S3E). Consistently, knockdown of ULK3 repressed accumulation of LC3C, but not LC3B, in response to CQ treatment in all cell lines (Figure 3A and 3B). In contrast, knockdown of ULK1 or combined knockdown of ULK1 and ULK2 inhibited CQ-dependent accumulation of LC3B, but not LC3C (Figure S3F, S3G, S3H). This implicates tissue specific activities of ULK3 in renal cells. Reexpression of exogenous Strawberry (STR)-tagged ULK3 to the same level as endogenous LC3C rescued LC3C flux (Figure 3C). STR-ULK3 colocalized with endogenous LC3C (Figure 3D) but not with endogenous LC3B (Figure 3E). The effects of ULK3 on LC3C did not involve changes in LC3C mRNA levels (Figure S3I). ULK3 was part of the complexes that include FIP200, ATG13 and LC3C (Figures 3F, 3G and 3H).

**Figure 3.**
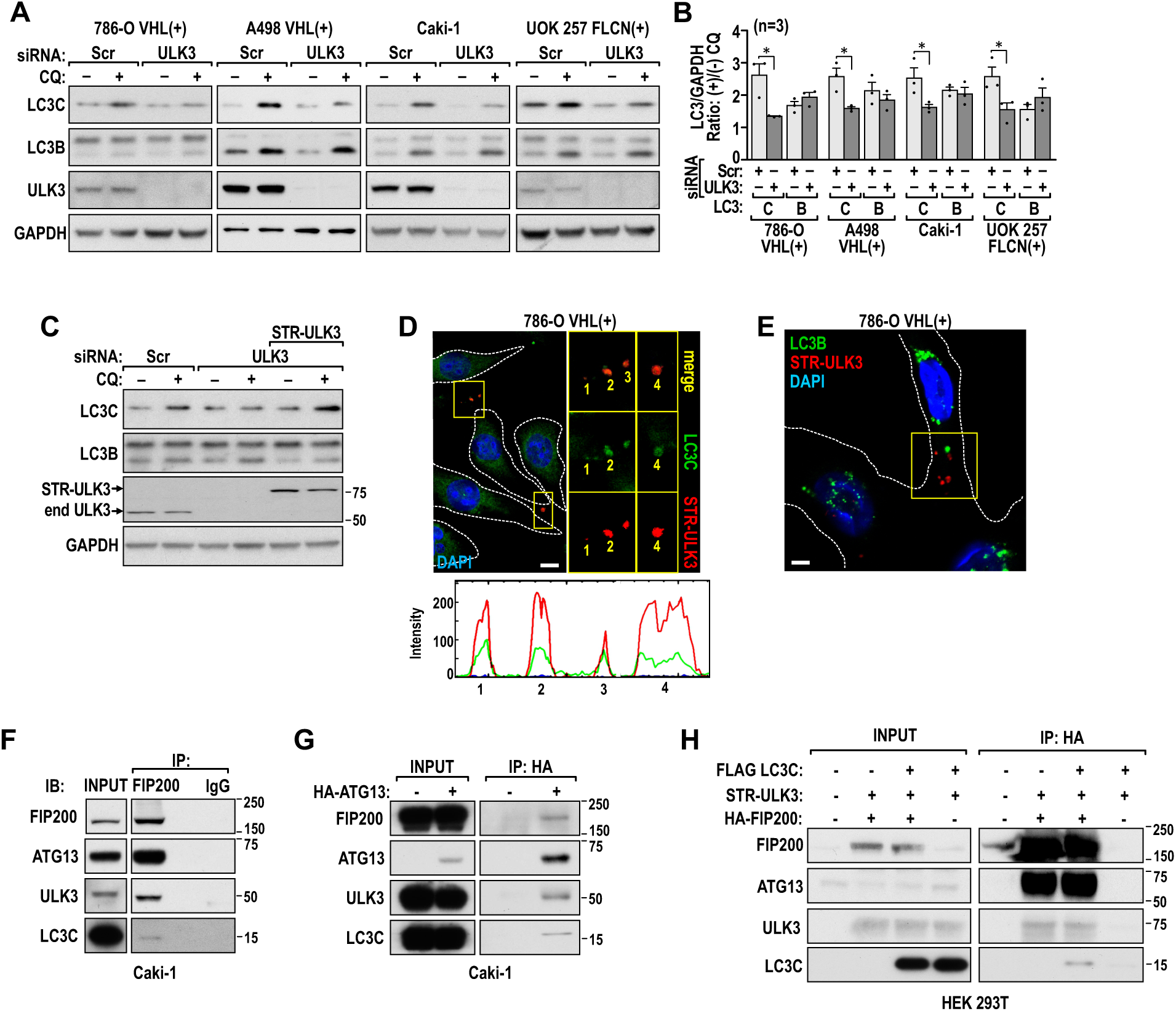
ULK3 is required for LC3C autophagy. (A) Immunoblots show decrease in the CQ dependent accumulation of LC3C in response to ULK3 knockdown in indicated cell lines. (B) Quantification of LC3C and B accumulation presented as the ratio of their protein levels normalized to GAPDH in CQ(+) vs. CQ(-) in cells transfected with scrambled or ULK3 siRNAs in the indicated cell lines. (C) Rescue of LC3C autophagic flux by reexpression of exogenous Strawberry-tagged ULK3 (STR-ULK3) to levels similar to that of endogenous ULK3 (end ULK3). (D) STR-ULK3 colocalizes with endogenous LC3C. Split channel RGB plots are shown. Cells were treated with 100 µM CQ for 1 hr. (E) STR-ULK3 does not colocalize with LC3B. (F) Co-immunoprecipitation of endogenous ULK3, ATG13 and LC3C with endogenous FIP200 in Caki-1 cells. (G) Co-immunoprecipitation of endogenous ULK3, FIP200 and LC3C in Caki-1 cells transfected with HA-ATG13. (H) Co-immunoprecipitation of ULK3, ATG13 and LC3C with FIP200 in HEK293 cells transfected with the indicated constructs. Statistical analysis as in Figure 1. Scale bars: 10 µm.

We also determined that ATG5 and ATG16 were necessary for both LC3C and LC3B lipidation (Figures S3J and S3K), and LC3C colocalized with ATG16L1 and ATG5 and FYVE puncta (Figure S3L, S3M), an indication that the lipidating cascade is required for LC3C autophagic program. Overall these data show that LC3C is an autophagic program regulated by a novel noncanonical upstream ULK- and PIK3C-associated complexes.

### LC3C autophagy targets PDMBs for degradation and prevents their asymmetric inheritance

Previous work showed that ULK3 localizes to the midbody and serves as an abscission checkpoint to assure full separation of DNA to the two daughter cells (Caballe et al., 2015). Consistently, we found that STR-ULK3 co-immunoprecipitates endogenous PDMB proteins MKLP1, CYK4, and CEP55 (Figure S4A). In addition, UVRAG localizes to midbodies (Thoresen et al., 2010). Thus, we hypothesized that LC3C, downstream from ULK3, regulates degradation of PDMBs during serum starvation. We found that under serum starvation conditions that induce LC3C flux (48 hr in 0.1% serum), LC3C, but not LC3B, colocalized with PDMBs (Figures 4A, S4B, 4B and 4C). Because LC3C is positively regulated by VHL and FLCN, we determined effects of VHL and FLCN on the accumulation of PDMBs. The number of PDMBs in *VHL*(-) or *FLCN*(-) RCC cells was high and did not increase by incubation with CQ for 24 hr (Figures 4D, 4E, 4F). Reconstitution of VHL in *VHL*(-) RCC cells or of FLCN in FLCN(-) cells lowered the number of PDMBs and recovered regulation by CQ, indicating the degradation of PDMBs by lysosomes (Figures 4D and 4E). In contrast, degradation of PDMBs in cells grown in 10% serum did not depend on *VHL* or *FLCN* status (Figures S4C and S4D), suggesting the existence of different autophagic programs degrading PDMBs under serum-starved and non-starved conditions. Importantly, knockdown of LC3C, but not LC3B, reversed the effects of VHL or FLCN reconstitutions, resulting in higher constitutive PDMB numbers and lack of regulation by 24 hr CQ treatments (Figures 4G and 4H). Knockdowns of LC3C-specific upstream regulators, ULK3, PIK3C2A, UVRAG, and RUBCN induced numbers of PDMBs similarly to the LC3C knockdowns supporting their role in degradation of PDMBs (Figure 4I). Consistently, there was colocalization of EGFP-2xFyve, a reporter for PI3P, with PDMBs markers that was significantly diminished by knockdown of PIK3C2A but not PIK3C3 (Figures 4J and 4K), further supporting the role PIK3C2A in the autophagic degradation of PDMBs by LC3C. Moreover, we found increased accumulation of the PDMB proteins MKLP1, CYK4, and ARF6 responsible for the integrity of PDMBs, in response to 24 hr CQ treatment in cells with intact LC3C. In cells with LC3C knocked down, these proteins accumulated at high levels in the absence of CQ, and there was no further induction in response to CQ treatment (Figures 4L and 4M). PDMBs colocalized with lysosomal marker LAMP1 (Figures 4N, S4E and 4O), further supporting their lysosomal degradation.

**Figure 4.**
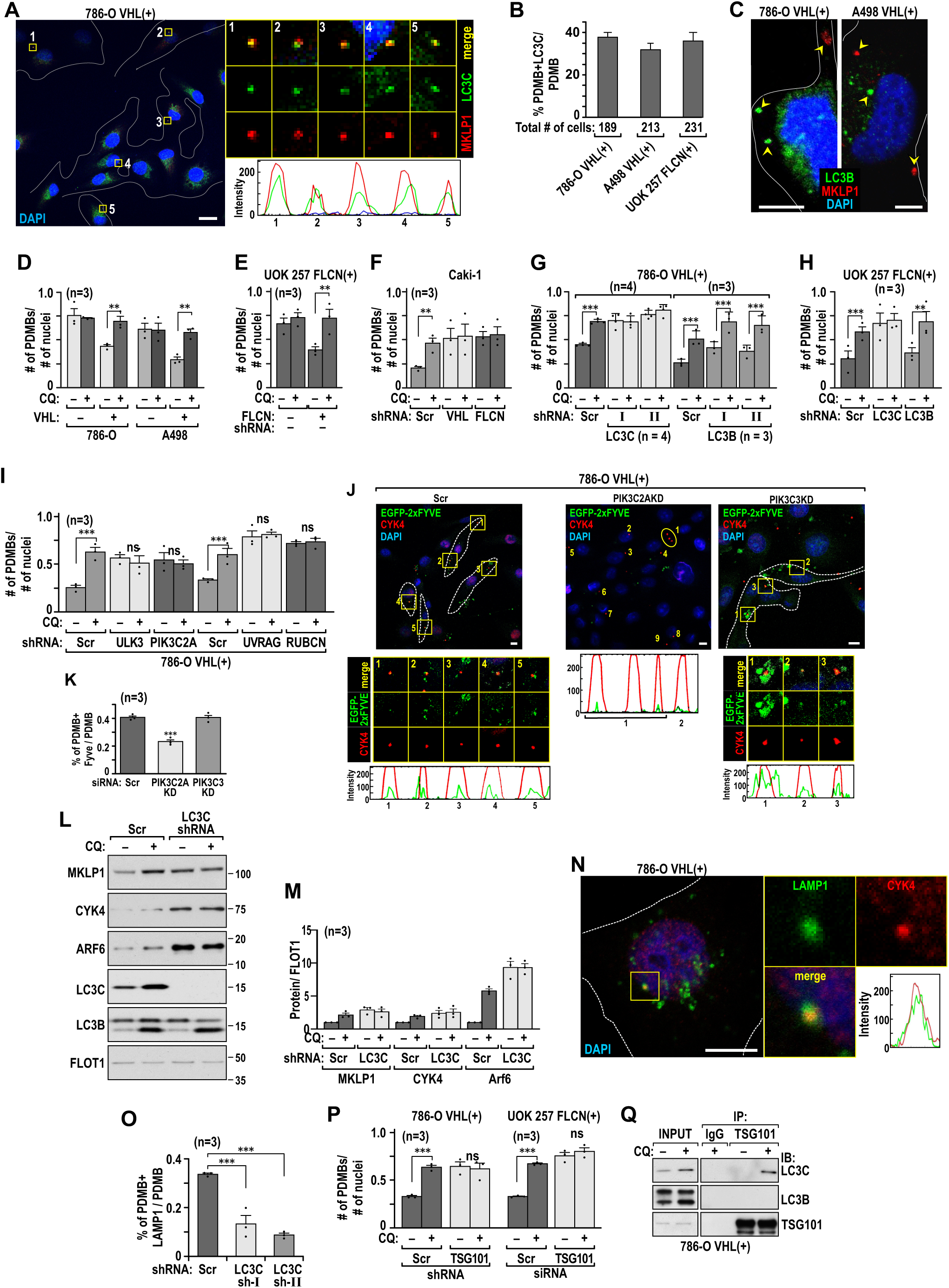
The LC3C autophagic pathway targets PDMBs for lysosomal degradation. (A) Endogenous LC3C colocalizes with PDMBs, labeled by MKLP antibody. Split channels and RGB plots are shown. Cells were treated with 100 µM CQ for 1 hr. (B) Quantification of the PDMBs colocalized with LC3C compared to the total number of PDMBs in indicated cell lines. (C) LC3B does not colocalize with PDMBs. Arrowheads point to the lack of colocalization of LC3B and PDMBs. Scale bar: 5 µm. (D) VHL represses the number of accumulating PDMBs and recovers CQ-dependent lysosomal degradation in cell lines without and with VHL. Ratios of the number of PDMBs to the number of nuclei are shown. Cells are treated 30 µM CQ for 24 hr. *n* is the number of independent experiments in which 10 fields with comparable numbers of cells were counted for PDMBs (labeled for MKLP1) and nuclei (DAPI). (E) Quantification of PDMBs in UOK 257 cell line without and with FLCN. (F) Quantification of PDMBs in Caki-1 cell line with knockdowns of VHL or FLCN. (G) Knockdown of LC3C but not LC3B induces number of PDMBs and abolishes CQ-dependent accumulation in 786-O VHL(+) cells. (H) Knockdown of LC3C but not LC3B induces PDMBs and abolishes CQ-dependent accumulation in UOK 257 FLCN(+) cells. (I) Knockdowns of LC3C upstream regulators, ULK3, PIK3C2A, UVRAG, and RUBCN consistently abolish CQ regulation of PDMBs and increases the number of PDMBs. (J) Colocalization of PDMB (CYK4) with PI3P reporter EGFP-2xFyve in cells with PIK3C2A and PIK3C3 knockdowns. (K) Quantification of PDMBs colocalized with EGFP-2xFyve in cells shown in (J). (L) Immunoblot shows CQ-dependent accumulation of PDMBs proteins, MKLP1, CYK4, and ARF6 in cells with LC3C and CQ-independent accumulation in cells with LC3C knockdown. (M) Quantification of immunoblots shown in (L) from three independent experiments. (N) Immunofluorescence experiment shows colocalization of PDMB with LAMP1 lysosomal protein. Split channels and RGB profiles are shown. (O) Quantification of colocalization PDMBs with LAMP1 in 786-O VHL(+) cells with and without LC3C. (P) Quantification of PDMBs in the indicated cell lines with TSG101 knockdowns. (Q) Endogenous TSG101 co-immunoprecipitates endogenous LC3C but not LC3B. Cells were treated with 100 µM CQ for 1 hr. Mean ± SEM are shown and P value is calculated by unpaired two-tailed t-test. If not otherwise indicated scale bars = 10 µm.

During cytokinesis, midbodies recruit the ESCRT (Endosomal Sorting Complex Required for Transport) complex. ULK3 interacts with the members of the ESCRT complex during cytokinesis (Caballe et al., 2015), thus we investigated role of ESCRT proteins in LC3C autophagy. We determined that the ESCRT-I member, TSG101, was necessary for the degradation of PDMBs (Figure 4P). Endogenous TSG101 co-immunoprecipitated endogenous LC3C but not LC3B (Figure 4R). Moreover, TSG101 colocalized with LC3C but not LC3B (Figures S4F, S4G). Knockdowns of TSG101 inhibited LC3C autophagy (Figure S4H) and mRNA (Figure S4J), pointing to direct and indirect effects of TSG101 on LC3C autophagy and expression.

The inheritance of the cytokinetic midbody from the most recent cell division can be asymmetric, if it is maintained by one of the daughter cells (Crowell et al., 2013; Kuo et al., 2011; Salzmann et al., 2014). The inheritance of PDMBs from earlier divisions is not known. Because LC3C degrades PDMBs, we hypothesized that LC3C would increase the number of symmetric divisions, where the PDMB is absent in either daughter cell, with only a new cytokinetic midbody present. Asymmetric division is defined as when one of the daughter cells maintains one or more PDMBs in addition to the cytokinetic midbody (Figure 5A). 786-O with lost *VHL* had approximately equal numbers of symmetric and asymmetric divisions (Figure 5B). In contrast, cells expressing reconstituted *VHL* had significantly higher numbers of symmetric vs. asymmetric divisions (Figure 5B). Knockdown of *VHL* in Caki-1 cells increased the number of asymmetric divisions (Figure 5C). Similarly, cells expressing FLCN had more symmetric divisions compared to cells without *FLCN* (Figure 5D). Knockdown of *LC3C* induced a significantly greater number of asymmetric divisions in cells expressing VHL or FLCN (Figures 5E, see also Figure7D). The LC3C-dependent clonal inheritance of PDMBs maintenance was further confirmed in cells grown as 3-dimensional spheroids. Cells expressing LC3C show low, while cells with LC3C knocked down show high numbers of inherited PDMBs (Figure 5F and 5G). Importantly, human ccRCCs with mutated *VHL* have higher numbers of PDMBs compared to tumors with wild type *VHL* (Figure 5H).

**Figure 5.**
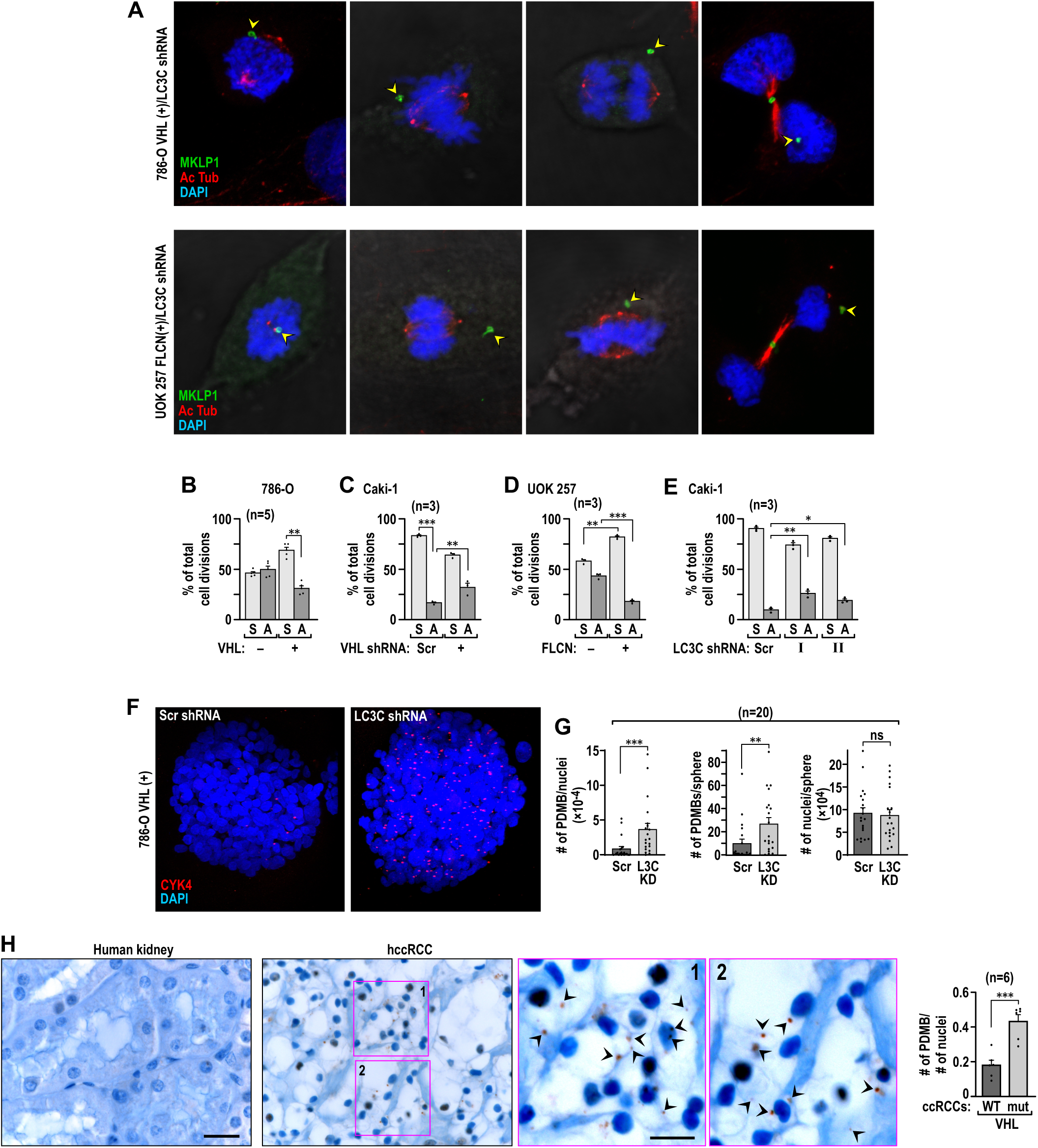
LC3C inhibits asymmetric inheritance of PDMBs. (A) Examples of asymmetric cell division in different phases of cell division in 786-O VHL(+) and UOK 257 FLCN(+) LC3C-knockdown cells. Arrowheads point to the PDMBs, which are different from the cytokinetic midbody in the intercellular bridge. Scale bar: 10 µM. (B) Percentage of symmetric (S) and asymmetric (AS) cells divisions compared to the total number of cells divisions in 786-O VHL(+) cells without and with reconstituted VHL. (C) Percentage of symmetric (S) and asymmetric (AS) cells divisions compared to the total number of cell divisions in Caki-1 cells with endogenous VHL and with VHL knocked down. (D) Percentage of symmetric (S) and asymmetric (AS) cells divisions compared to the total number of cells divisions in UOK 257 cells without and with FLCN. (E) Knockdown of LC3C increases percentages of asymmetric (AS) vs. symmetric (S) cell divisions as compared to the total number of cells divisions in Caki-1 cells with LC3C knocked down. (F) 786-O VHL(+) cells with LC3C knockdown grown as 3-dimensional spheroids maintain higher levels of PDMBs (CYK staining) in multiple daughter cells. (G) Quantification of PDMBs, nuclei, and ratios of PDMBs/nuclei for 20 spheres in each category. Scale bar: 20 µM. (H) Immunocytochemistry for PDMBs (CYK4 staining, arrowheads point to PDMBs) and quantification of PDMBs in sections from human normal kidney and ccRCCs with wild type or mutated *VHL*. Magnified inserts are shown to the right. Mean ± SEM are shown; P value is calculated by unpaired two tailed t-test.

Overall these data provide evidence that the fate of PDMBs, markers of stemness, is under tight control by renal tumors suppressors VHL and FLCN and their downstream target, the LC3C autophagic program.

### LC3C autophagy requires the LIR motif on CALCOCO2 and the LIR-binding motif on LC3C

Two major differences between LC3C and LC3B are the CLIR-binding motif in addition to the LIR-binding motif in the N-terminal region and the presence of the C-terminal peptide. Selectivity of LC3C autophagy towards bacterial pathogens requires interaction of LC3C with the CALCOCO2 cargo receptor in a manner depending on the protein interaction between a specific CLIR motif on CALCOCO2 and a CLIR-binding motif in the N-terminal region of LC3C (von Muhlinen et al., 2012). However, LC3C also contains a LIR-binding site. Thus, we investigated the role of CALCOCO2 and LIR and CLIR motifs in LC3C autophagy. Knockdowns of CALCOCO2, p62, and NBR1 (Figure S5A) in cells expressing LC3C demonstrated that only knockdown of CALCOCO2 abolished PDMB degradation, while knockdowns of the other two cargo receptors were without effect (Figure 6A). CALCOCO2 colocalized with LC3C (Figures 6B) and with PDMBs (Figures 6C, 6D and S5B). This further supports the idea that LC3C-dependent PDMB degradation is a program independent from the previously described autophagic degradation of midbodies, which requires p62 or NBR1 (Kuo et al., 2011; Pohl and Jentsch, 2009). LC3C with a mutated CLIR-binding motif (Figure S5C) rescued lysosomal degradation of PDMBs (Figure 6E). Consistently, CALCOCO2 with a mutated CLIR motif (Figure S5D), reconstituted in cells with CALCOCO2 knocked down, rescued PDMB lysosomal degradation similarly to the wild type CALCOCO2 (Figure 6F). In contrast, CALCOCO2 with a mutated LIR motif (Figure S5D) failed to rescue PDMB lysosomal degradation (Figure 6G). This indicates that CALCOO2 may interact with LC3C through either the LIR or CLIR (von Muhlinen et al., 2012) motif, most likely depending on the biological context of selective degradation.

**Figure 6.**
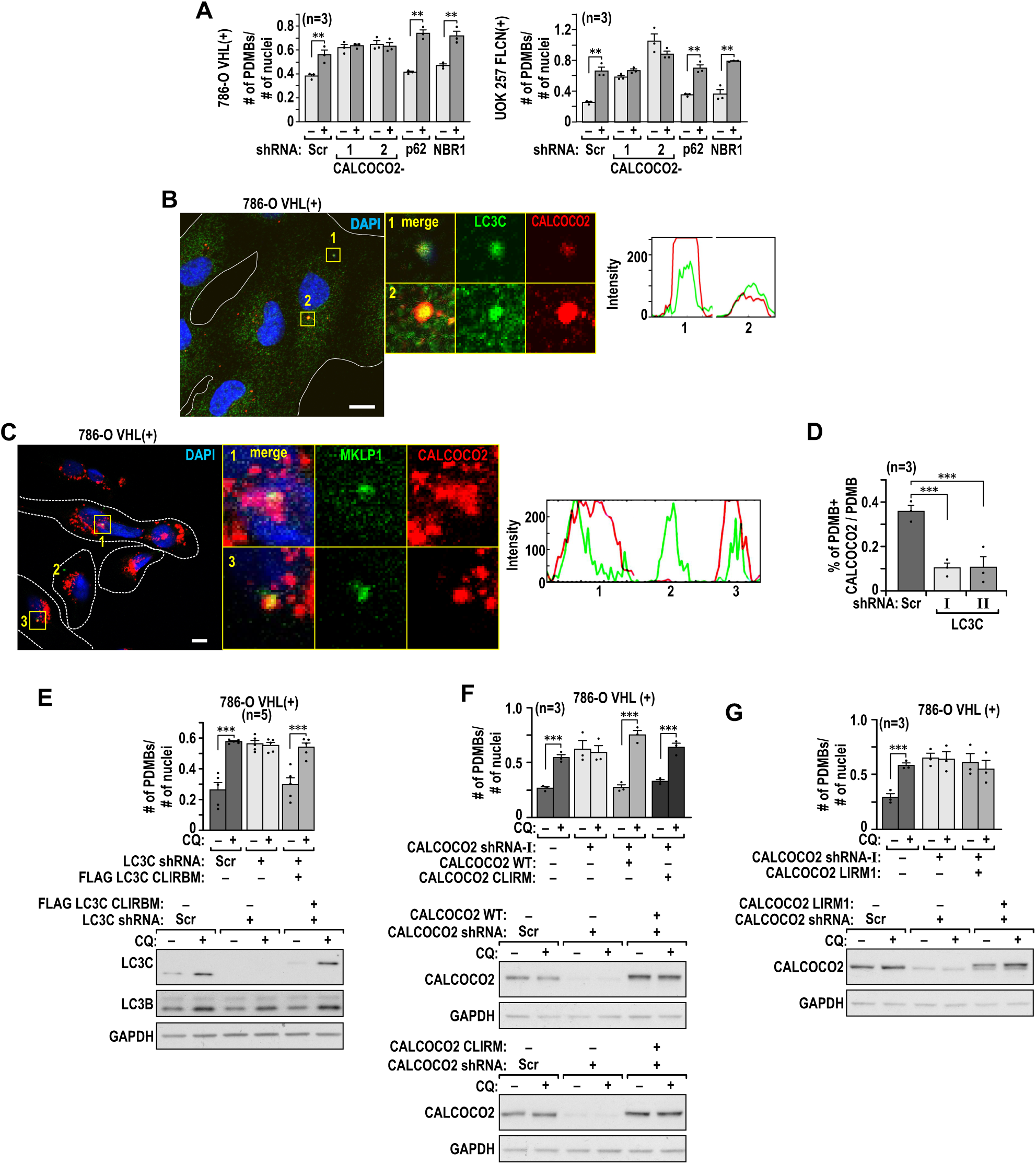
LC3C autophagy requires CALCOCO2 cargo receptor and LIR motif. (A) CALCOCO2 cargo receptor, but not p62 or NBR1, is necessary for the degradation of PDMBs by lysosome in the indicated cell lines. (B) Colocalization of endogenous LC3C with endogenous CALCOCO2. RGB profiles and split channels are shown. (C) CALCOCO2 colocalizes with PDMB. Split channels and RGB profiles are shown. (D) Quantification of PDMBs colocalized with CALCOCO2 in cells with and without LC3C. (E) Flag-tagged mutant of CLIR-binding site on LC3C (CLIRBM) is able to rescue CQ-dependent degradation of PDMBs in cells with LC3C knocked down. Immunoblot shows expression of FLAG-tagged LC3C-CLIRBM at the level of endogenous LC3C. (F) CLIR mutant of CALCOCO2 (CLIRM) is able to rescue CQ-dependent degradation of LC3C in cells with CALCOCO2 knocked down, similarly to wild type CALCOCO2. Immunoblots show expression of exogenous CALCOCO2 wild type and CLIRM matched to the expression of the endogenous CALCOCO2. (G) LIR mutant of CALCOCO2 is not able to rescue CQ-dependent degradation of PDMBs in cells with CALCOCO2 knocked down. Immunoblot shows expression of exogenous CALCOCO2 LIRM matched to the expression of the endogenous CALCOCO2. Analysis of PDMBs was performed as described in Figure 4D. Means± SEM are shown. P value is calculated by unpaired two tailed t-test Scale bars: 10 µM.

### LC3C C-terminal peptide is necessary and sufficient for selectivity of LC3C autophagy towards PDMBs

Here we investigated the role of the C-terminal peptide in selectivity of LC3C autophagy. Reexpression of the autophagy-deficient LC3C G126A mutant (Figure S6A) in cells with LC3C knocked down did not rescue lysosomal degradation of PDMBs, further supporting the role of LC3C autophagy in regulation of PDMB fate (Figures 7A). Reconstitution of the wild-type LC3C, but not its short form without the C-terminal peptide (C127Stop) (Figure S6A), rescued lysosomal degradation of PDMBs (Figures 7B and 7C) and the number of symmetric cells divisions (Figure 7D). This indicates that the C-terminal peptide is necessary for LC3C-degradation of PDMBs. Interestingly, accumulation of the LC3C C127Stop protein was induced by lysosomal inhibition with CQ. This means that LC3C without the peptide is active in autophagy, but likely targets different cargo than the full size LC3C. In order to determine if the C-terminal peptide is sufficient for degradation of PDMBs, we generated a chimeric LC3B-C construct, where the C-terminal peptide was added after G120 in LC3B (Figure S6B). This construct was expressed to levels similar to the expression of endogenous LC3C as measured by RT PCR with primers selective for LC3C and LC3B-C (Figure S6C). The chimera was able to restore regulation of PDMBs similarly to what was measured in cells expressing LC3C (Figure 7E). Moreover, in cells expressing the chimera, PDMB colocalized with LC3B or were in close proximity (Figure 7F and S6D).

**Figure 7.**
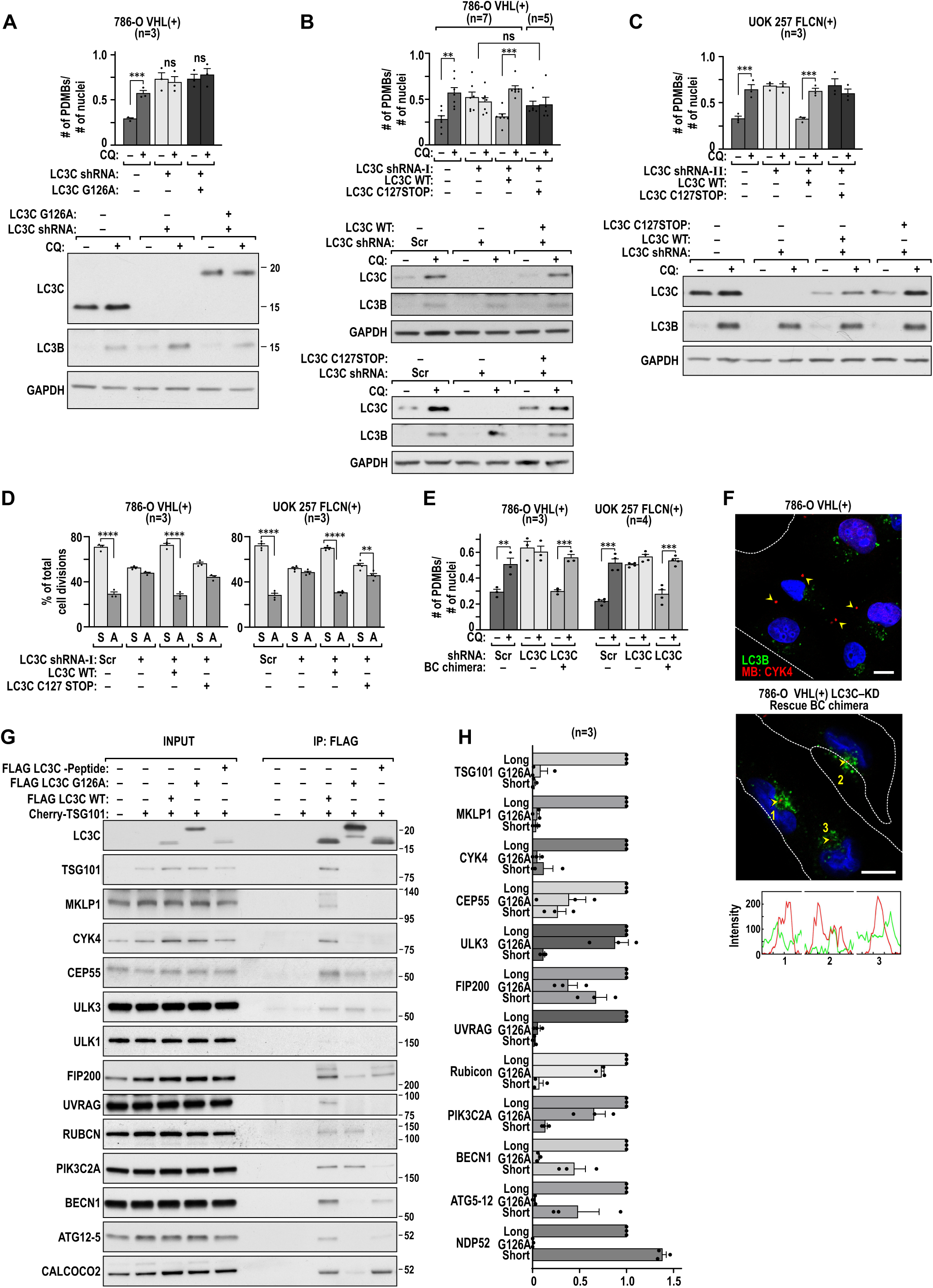
LC3C C-terminal peptide is necessary and sufficient for selectivity of LC3C autophagy towards PDMBs. (A) Expression of autophagy defective mutant of LC3C, G126A, in cells with LC3C knocked down does not rescue PDMBs degradation. Immunoblot shows expression of LC3C mutant. (B) Reconstitution of LC3C wild type, but not the C127Stop without the LC3C peptide, in 786-O VHL(+) cells rescues PDMB degradation. Immunoblots show levels of expression of exogenous LC3C matches levels of endogenous LC3C. (C) Reconstitution of LC3C wild type, but not the C127Stop without the LC3C peptide, in UOK 257 FLCN(+) cells rescues PDMBs degradation. Immunoblots show levels of expression of exogenous LC3C matches levels of endogenous LC3C. (D) Expression of wild type LC3C, but not the C127stop mutant, rescues the number of symmetric divisions in the indicted cell lines. (E) Expression of LC3B-LC3C C-terminal peptide chimera rescues regulation of PDMBs by lysosomal degradation. (F) Representative examples of immunofluorescence experiments showing lack of colocalization of PDMBs with LC3B in cells with LC3C knockdown (left), but appearance of close proximity and colocalization in cells expressing BC chimeras (right). RGB profiles are shown. (G) Co-immunoprecipitation of FLAG-tagged LC3C wild type and C127Stop mutants with endogenous PDMBs proteins in the presence of co-expressed Cherry-TSG101. Experiments were performed in HEK293T cells. (H) Quantification of the experiments shown in (G). Mean ± SEM are shown; P value is calculated by unpaired two tailed t-test. If not otherwise indicated scale bars = 10 µm.

Next, we sought to determine if binding of PDMB proteins and formation of the noncanonical complex with LC3C was regulated by the presence of LC3C peptide. We expressed flag-tagged LC3C wild type and C127Stop mutant together with TSG101 in HEK293T cells and determined binding of endogenous PDMBs proteins and LC3C autophagic regulators by co-immunoprecipitation. Only wild-type LC3C co-immunoprecipitated TSG101, all three PDMB markers, and members of the noncanonical upstream complexes ULK3, UVRAG, RUBCN, and PIK3C2A (Figures 7G and 7H). Moreover, the canonical regulators of autophagy, FIP200, BECN1, and ATG5-12 conjugate co-immunoprecipitated with both the long and short form of LC3C, but binding was stronger in the case of LC3C with C-terminal peptide (Figures 7G and 7H). Interestingly, CALCOCO2 bound similarly to the full size and short form of LC3C, indicating that interaction between LC3C and CALCOCO2 is not a major determinant of LC3C selectivity towards PDMBs. Importantly, some upstream regulators, such as ULK3, FIP200, Riubicon and PIK3C2A bound also to the autophagy deficient G126A LC3C mutant. This points to a potential sequence in the assembly of the upstream regulatory complexes before processing of LC3C by cleavage and lipidation. Overall the data indicate that LC3C C-terminal peptide is an importnat tool of functional selectivity of LC3C autophagy towards PDMBs and is necessary for the assembly of the noncanonical autophagic regulatory complex.

## Discussion

Selectivity in the lysosomal degradation of specific cargo under particular biological conditions is essential for maintenance of cellular homeostasis. Our data point to a complex plasticity in LC3C autophagic function that evolved in human cells versus other species. We identified a novel mechanism of selectivity in LC3C autophagy that requires its C-terminal peptide. However, we also noticed that LC3C mutant without the C-terminal peptide, that does not degrade PDMB cargo, accumulates in response to lysosomal inhibition, an indication that this short form of LC3C is engaged in autophagy as well. Moreover, LC3C has ability to interact with cargo receptors through LIR- or/and CLIR-binding motifs. This indicates that LC3C can participate in diverse autophagic processes, selectively mediated by different utilization of LIR or CLIR binding motifs, and absence or presence of the peptide.

We propose that LC3C represents a novel hybrid autophagic program at the intersection of classic autophagy and the endo/phagolysosomal pathway. Clearly, in all four renal cancer derived cell lines LC3C required activity of ULK3, rather than ULK1/2, in the complex associated with FIP200/ATG13. The activity of LC3C-dependent autophagy requiring ULK1 has been reported in the case of xenophagy, however the role of LC3C peptide was not evaluated in these experiments (Ravenhill et al., 2019). Together, these findings further support the role of LC3C as a multifaceted regulator that can require either ULK3- or ULK1-associated complex depending on the biological context.

We have initially determined that LC3C autophagy required Beclin 1 and the E3-like ATG5/12/16 lipidating cascade. There are two Beclin 1-associated autophagosome initiating complexes. Both contain PIK3C3, but complex I includes ATG14, while complex II incorporates UVRAG but not ATG14 (Liang et al., 2006). The activity of Beclin 1-UVRAG complex is inhibited by Rubicon (Matsunaga et al., 2009). UVRAG has been shown to stimulate both autophagosome and endosome maturation promoting lysosomal degradation of cargo, independently from its role in autophagosome initiation (Liang et al., 2008). The LC3C requirement for UVRAG supports the role of Beclin 1-complex II activity; however, surprisingly, Rubicon is a positive regulator of LC3C. This points towards similarity to the recently described Beclin 1/UVRAG/PIK3C3/LC3B-dependent pathway of LC3-associated phagocytosis, i.e. LAP, where Rubicon is a positive regulator (Lai and Devenish, 2012; Martinez et al., 2016; Martinez et al., 2015; Sanjuan et al., 2007). LAP is not considered a macroautophagic program because it involves formation of a single membrane structure originating at the plasma membrane to sequester extracellular pathogens and cell debris. It does not require ULK1 but shares several other regulators with autophagy, such as ATG3, ATG5, ATG7, ATG12, and ATG16, which ultimately leads to recruitment of lipidated LC3s on the intravesicular side only, thus preventing recycling of LC3. Pathogens are recognized by plasma membrane receptors and there is no evidence for ubiquitin modifications. LC3C participates in selective autophagy (xenophagy) of S. Typhimurium in a mechanism requiring CALCOCO2 and the CLIR motif on CALCOCO2 (von Muhlinen et al., 2012). In contrast to LAP, xenophagy requires formation of a double membrane autophagosome around a free bacterium in the cytosol, using standard pre- and initiating complexes. The intracellular pathogens are ubiquitylated. However, LAP and xenophagy may have more in common, eg. fusion of phagosomes with autophagosomes, or recruitment of phagosomes as the source of the autophagosome initiating membrane. Potential involvement of LC3C with a LAP-like program in degradation of PDMBs is supported by the fact that midbodies released during cell division can undergo actin-dependent phagocytosis (Crowell et al., 2014; Fazeli et al., 2016).

The fact that LC3C function requires PIK3C2A and TSG101 further supports LC3C function at the intersection between autophagosome and endosome. Previous work indicated that knockdowns of PIK3C2A decreased autophagic output and accumulation of recycling endosomes (Merrill et al., 2017). TSG101 is a member of ESCRT complex used by the endolysosomal pathway that delivers cargo originating at the plasma membrane or from the extracellular environment via multivesicular bodies (MVBs). Biogenesis of MVBs requires a cascade of events regulated by ESCRT to form large numbers of small intraluminal vesicles (IVLs) from the limiting membrane that sort the cargos for lysosomal degradation.

Consistent with the tumor suppressing activity of LC3C, Beclin 1 and UVRAG are tumor suppressors and were shown to cooperate to induce autophagy and suppress tumorigenicity of colon cancer cells (Liang et al., 2006). TSG101 and PIK3C2A are favorable prognostic factors in ccRCC according to The Human Protein Atlas. LC3C tumor suppressing activity targets Met RTK and regulates cell migration and invasion (Bell et al., 2019). It remains to be determined if LC3C autophagy can have both tumor suppressing and oncogenic activities, depending on the presence or absence of the C-terminal peptide. We established that LC3B has oncogenic activity in ccRCC (Hall et al., 2014; Mikhaylova et al., 2012), thus expression of short form of LC3C may mimic effects of LC3B.

LC3C-dependent autophagic degradation of PDMBs occurs under conditions of low concentration of serum, which is different from previously reported autophagic degradation of cytokinetic midbodies performed with standard cell culture media containing 10% serum. In those studies, p62 and NBR1 were identified as cargo receptors (Kuo et al., 2011; Pohl and Jentsch, 2009). Moreover, these studies investigated the inheritance of midbodies formed during the most recent cytokinesis (first generation). We, however, investigated inheritance of PDMBs, i.e. midbodies that were formed during previous cytokinesis. Importantly, the maintenance of PDMBs was heritable; clonal spheroids were easily separated into those with high and low PDMB content according to LC3C status. This implies that regulation of PDMBs by LC3C contributes to cell fate reprogramming.

We have demonstrated that the C-terminal peptide is functionally necessary for the degradation of PDMBs and for the assembly of the LC3C regulatory complexes. Interestingly, we see exclusively the lipidated form of LC3C that accumulates in response to lysosomal inhibitors and never the noncleaved form, which indicates very fast processing of LC3C to its active form. Recent evidence was presented for an interaction of TBK1-phosphorylated serines 93 and 96 in the N-terminal region of LC3C with arginine 134 in the peptide, promoting local structuring of the otherwise highly dynamic and unstructured peptide. This interaction diminished ability of ATG4 to cleave the peptide (Herhaus et al., 2020). These data support the functional and regulatory role of the peptide in LC3C autophagy. We have observed robust transcriptional regulation of LC3C by serum starvation, a condition that induces LC3C autophagy in multiple cell lines (Figure S1 (Mikhaylova et al., 2012). Strong dependence on the continuing input of new LC3C molecules is likely to be necessary in the LC3C autophagy requiring C-terminal peptide, as only the newly translated LC3C will have the peptide, while LC3C that is cleaved of the preexisting autophagosome and reused will not, eliminating functional activity of the peptide. Mechanistic understanding of the regulatory role of the C-terminal peptide will be subject of future studies.

## Material and methods

### Cell lines and basic protocol

Human isogenic VHL(-) and VHL(+) 786-O, A498 cells, Caki-1 cells with endogenous wild type VHL and FLCN and their knockdowns, and HEK293T were previously described (Bastola et al., 2013; Hall et al., 2014; Mikhaylova et al., 2012). UOK 257 cells have inactivated FLCN. Isogenic cell lines with reconstituted FLCN were a gift from Dr. Laura Schmidt (NCI). All cells were grown in DMEM-F12 (Hyclone SH30023) with 10% FBS. Human renal proximal tubule epithelial (TH1) cell line was grown in DMEM-High glucose media (Hyclone SH30022) with 10% FBS according to supplier’s recommendations. All cell lines are periodically authenticated (Genetica, Cincinnati, OH) and media is monthly checked for Mycoplasma. All relevant reagents are listed in Table S1.

Unless otherwise indicated, experiments were performed with the following timeline: cells (10-30 x10^4^) were plated in 60 mm plates in DMEM/F12 medium with 10% serum and allowed to attach. Medium was changed to contain 0.1% serum and cells were collected 48 hr later. Specific treatments were performed at the end of the 48 hr period. For autophagic flux experiments cells were treated with 100 µM CQ or 200 nM Bafilomycin A1 for 1-4 hr before collection, and for counting PDMBs cells were treated with 30 µM CQ for 24 hr. RNA extractions and QRT-PCR were described (Hall et al., 2014; Mikhaylova et al., 2012). For determinations of symmetric and asymmetric cell divisions cells were placed in 0.1% starvation media as described above, chased with 1% serum in the final 12 to16 hr before collection. All antibodies, primers, and chemicals used are listed in Table S1.

### Transfections and transductions

RCC cell lines were transfected with siRNA and plasmids using Lipofectamine 3000 according to manufacturer’s protocol. HEK293T cells were transfected using 2 µg polyethylenimine per 1 µg DNA. SiRNAs were used at final concentrations of 50 nM or 80 nM. HEK293T cells were transfected with the following amounts of DNA: 100-500 ng of LC3C, 0.5 µg VHL, 2-5 µg Cherry-TSG101, 2.5 µg GFP-MKLP1, or 0.5 µg STR-ULK3. Viral transductions included addition of 1.1 µg/mL polybrene before viruses were administered. All lentiviral shRNA constructs were VSV-G envelope packaged and ten-fold concentrated (Cincinnati Children’s Hospital Medical Center Viral Vector Core), and used at a 1:30 dilution. All controls were treated with empty, non-target, or scrambled constructs. Media was changed 6-8 hr after transfection or the following day after transfection/transduction unless specified. Stable cell line pools were selected with appropriate antibiotics 2-3 days post transduction and maintained in the selection media.

To achieve similar expression levels compared to endogenous the following approaches were used: For UVRAG rescue experiments, knockdown of endogenous UVRAG was performed on cells stably expressing STR-UVRAG. For ULK3 and RUBCN rescue experiments cells were transduced with viral exogenous constructs at 1:5,000 and 1:20,000 respectively, then replated 3 days later for knockdown of the endogenous genes. For PIK3C2A, cells were transfected with 15 µg plasmid PIK3C2A DNA and resplit 3 days later, following knockdown of endogenous gene. For FLAG-LC3C rescue experiments 50-150 ng plasmid DNA was transfected in stable LC3C knockdowns. For CALCOCO2 reconstitutions, 0.3-0.4 µg DNA was transfected, media was changed to starvation 6-8 hours later and cells were treated with lentiviral packaged shRNA in the starvation media. In all cases cells were starved with 0.1% FBS for 48 hr before collection. For LC3C rescue with untagged viral constructs, virus was titrated between 1:1000-1:500,000 stably or transiently. For immunofluorescence 350-850 ng GFP-RUBCN, 250-500 ng GFP/FYVE, or 350-700 ng STR-ULK3 was transfected per well.

### Co-immunoprecipitations and immunoblotting experiments

For immunoblot analysis cells were collected in RIPA buffer (25mM Tris HCl pH 7.6, 150mM NaCl, 1% NP-40, 1% sodium deoxycholate, 0.1% SDS). 10-35 µg of extracts were separated on 3-8%, 12%, or 4-12% polyacrylamide gels and transferred onto PVDF membrane. Blots were probed with relevant antibodies, as listed in Table S1. Density of bands was quantified using Image J and normalized to appropriate loading control.

For immunoprecipitations, cells were lysed in buffer containing 50 mM HEPES pH 7.9, 5 mM MgCl_2_, 150 mM NaCl, 1% glycerol, 1% IGEPAL, or a buffer containing 50mM Tris pH 8.0, 137mM NaCl, 1mM MgCl_2_, 1mM CaCl_2_, 1% NP-40 and protein lysates were incubated with primary antibody in the same buffer with detergent concentration adjusted to 0.2%. 0.5-1 mg of cell lysates in 0.5-1 ml of indicated buffers supplemented with protease/phosphatase were incubated with primary antibodies overnight at 4°C, followed by 1 hr incubation with protein A/G or protein G magnetic beads at room temperature. In the case DynaBeads were used, the beads were incubate with the antibody first following with 10 min incubation with cell lysates. Beads were washed in the IP buffer, and co-immunoprecipitated proteins were eluted with 1x LDS sample buffer for 10 min at 45°C. In the cases where primary antibodies used for immunoblotting were from the same species as the antibody used for immunoprecipitations, Clean-Blot IP Detection Reagent (HRP) was used in place of a secondary antibody. For each immunoprecipitation reaction the following amounts of primary antibodies were used: 2 µg of anti-TSG101, 3 µg of anti-HA, 1:200 dilution of anti-Strawberry, either 2 µg or 1:250 of anti-FLAG M2, 1:100 dilution of anti-FIP200, 1:50 dilution of anti-UVRAG. The amount of control IgGs was adjusted to the concentrations of the primary antibodies.

### Immunofluorescence and PDMB count

For immunofluorescence analysis, cells plated on glass coverslips were fixed with 4% paraformaldehyde for 20 min at room temperature or 100% methanol for 5 min at −20°C. PFA fixed cells were additionally quenched with 50 mM NH_4_Cl unless already expressing a fluorescent marker. Cells were permeabilized with 0.3% Triton or 0.1% saponin. Coverslips were blocked with PBS containing either 0.1% Triton or 0.1% saponin and either 3% NGS or 1% BSA for 30 min and incubated with primary antibody for 1 hr at 37°C. Coverslips were then washed and incubated with Alexa Fluor labeled secondary antibodies for 30 min at room temperature. Finally, coverslips were washed and mounted using DAPI Fluoromount-G. Antibodies and concentrations are listed in Table S1. PDMB were counted based on labeling with anti-MKLP1 or anti-CYK4 antibodies. For each experiment, 10 independent and random fields of view were counted for the numbers of PDMBs and nuclei. Each section had between 10-30 cells. Ratios of number of PDMBs to number of nuclei from all sections were averaged. Confocal images were acquired on a Zeiss LSM710 confocal microscope with a Zeiss Axio.Observer Z1 stand and a Zeiss Plan-Apochromate objective (63x/1.4 Oil DIC) using the Zeiss Zen2010 software. The appropriate lasers and emission filters for the respective fluorophores were used in a multitracking mode. Widefield images were acquired using a Zeiss Axioplan 2 imaging microscope with the appropriate filter cubes and a Zeiss AxioCam MRm B&W camera to record the images using the Zeiss AxioViosion (Rel4.7) software. Used objective had the following specifications: Plan Apochromat 10/0.45; Plan Neofluar 20x/0.5; Plan Neofluar 40x/0.6 (air); Plan Neofluar 40x/1.3 (Oil); Plan Apochromat 63x/a.4 (Oil): alpha-PlanFluar 100x/1.45. All images were saved as 8-bit images.

Images were also acquired on a Nikon SIMe microscope with a Nikon Eclipse Ti stand using a Hamamatsu C11440 Orca Flash 4.0 camera and a Nikon SR Apo TIRF 100x SR/1.49 objective. DAPI images of nuclei were acquired as widefield images while we used the structured illumination capabilities (SIM) for the fluorescent channels with a 488nm, 561nm and 640nm laser line excitation.

RGB profiles (generated by ImageJ) and split channels are shown. For determination of symmetric vs. asymmetric cell divisions, coverslips were incubated with antibody against MKLP1 to visualize PDMBs and acetylated tubulin to visualize microtubules, and divisions were counted manually under the confocal microscope. For immunocytochemistry sections of fixed and paraffin-embedded tumors were processed in the Pathology Research Core at CCHMC and analyzed by light microscopy.

### Growth of cells as spheroids

Cells were grown in 8- or 16-well glass chamber slides using a dual chamber coat/culture layer. The floor of each chamber was coated with 100% liquid Matrigel (30-40 µL for 8 well slides and 20 µL for 16 well) and allow to solidify for 30 min in the tissue culture incubator. The culture layer was a 1:1 volume mixture of Matrigel and cells in the plating media (DMEM/F12 and 10% serum). 600 cells were plated in 8 well and 200 cells in 16 well slides. After addition of cells, Matrigel was allowed to solidify and media was added. The next day, media was replaced with fresh plating media. After four days, media was changed to the same media containing 0.1% serum. Spheroids were collected after 7-8 days from the time media was changed. For immunofluorescence analysis, spheroids were fixed using 100% methanol for 12 min at −20°C. Spheroids were permeabilized using 0.25% saponin in PBS for 10 min. Spheroids were rinsed with PBS, blocked for 1.5 hr using IF buffer (PBS, 0.1% saponin, 0.05% T-20, 0.05% NaN_3_) with 1% BSA at room temperature and incubated overnight with primary antibodies in IF buffer at 4°C. Incubation with secondary antibody in the IF buffer was for 1 hr at room temperature. Spheroids were washed and mounted using DAPI Fluoramount-G and analyzed by confocal microscopy.

### Statistical analysis

Data are expressed as mean ± SEM for independent experiments ≥ 3, or as mean ± SD for n=2. Analysis of differential expression was performed using unpaired two tailed t-test. *P<0.05; **P<0.01, ***P<0.001, ****P<0.0001

## ACKNOWLEDGMENTS

The work was supported by the following grants: R01CA122346, R01GM128216, 2I01BX001110 BLR&D VA Merit to M.F. C-K. J.M. was supported by UC P30-ES006096 CEG, R01MH107487, and R01DK091566. D.R.P was supported by R01CA168815. JLG was supported by R01NS094144 and R01CA211066. Y.Z. was partially supported by Oversea Study Stipend from Shandong University, Jinan, People’s Republic of China. We thank Dr. C. Wang for reading the manuscript, B. Peace for professional editing, and G. Doerman for preparing the figures.

## AUTHOR CONTRIBUTIONS

Conceptualization: M.F.C-K, J.M.; experiments and data collection: M.E.B., Y. Z, J.C., A.P., N.T. A.P, M.N.; confocal analysis: B.E.; discussions and reagents: J.L.G., D.R.P.; Y. Z.. Manuscript preparation: M.F.C-K, M.E.B., D.R.P, J.L.G.

## Supplemental Data

**Fig. S1.**
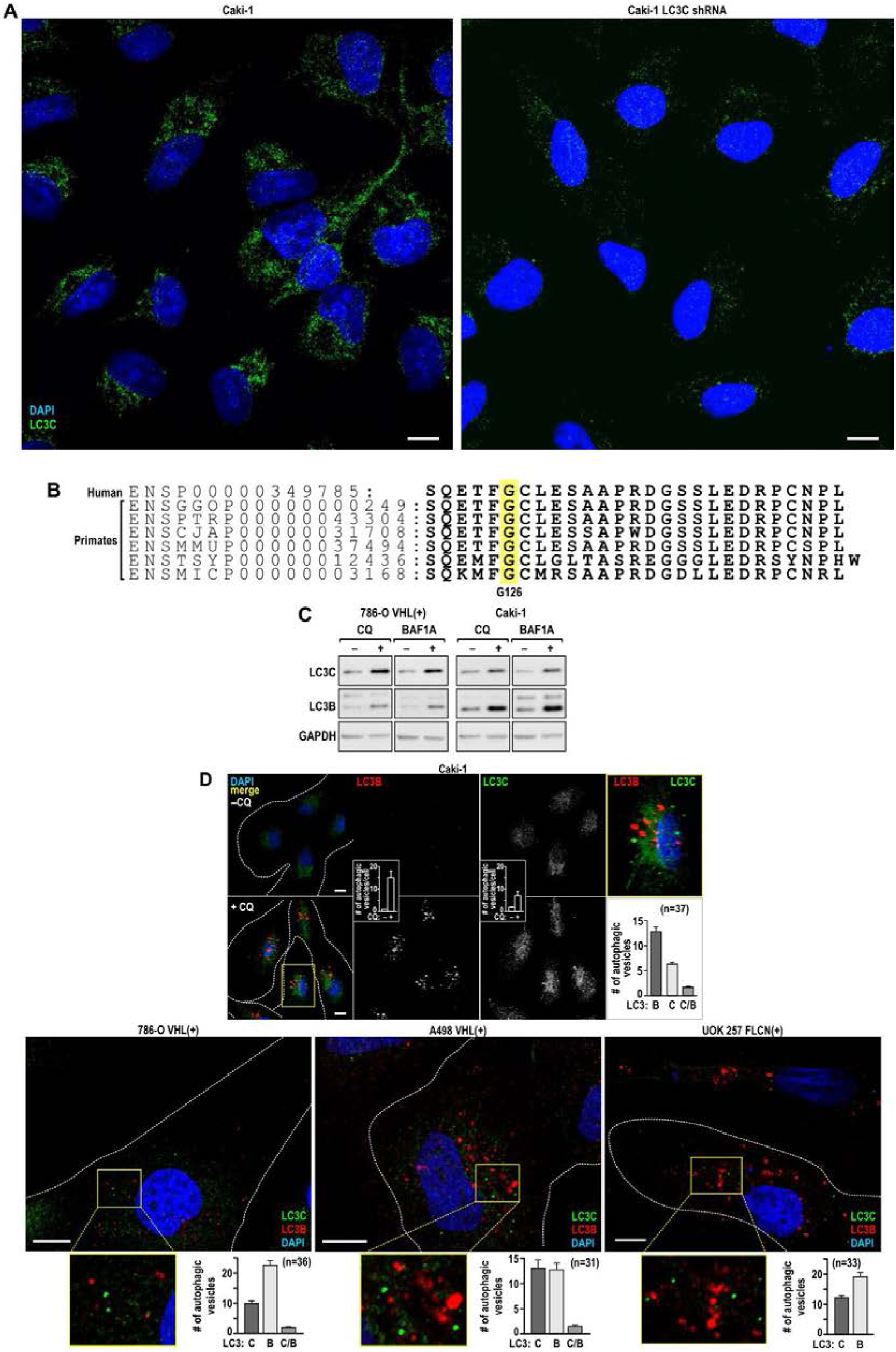
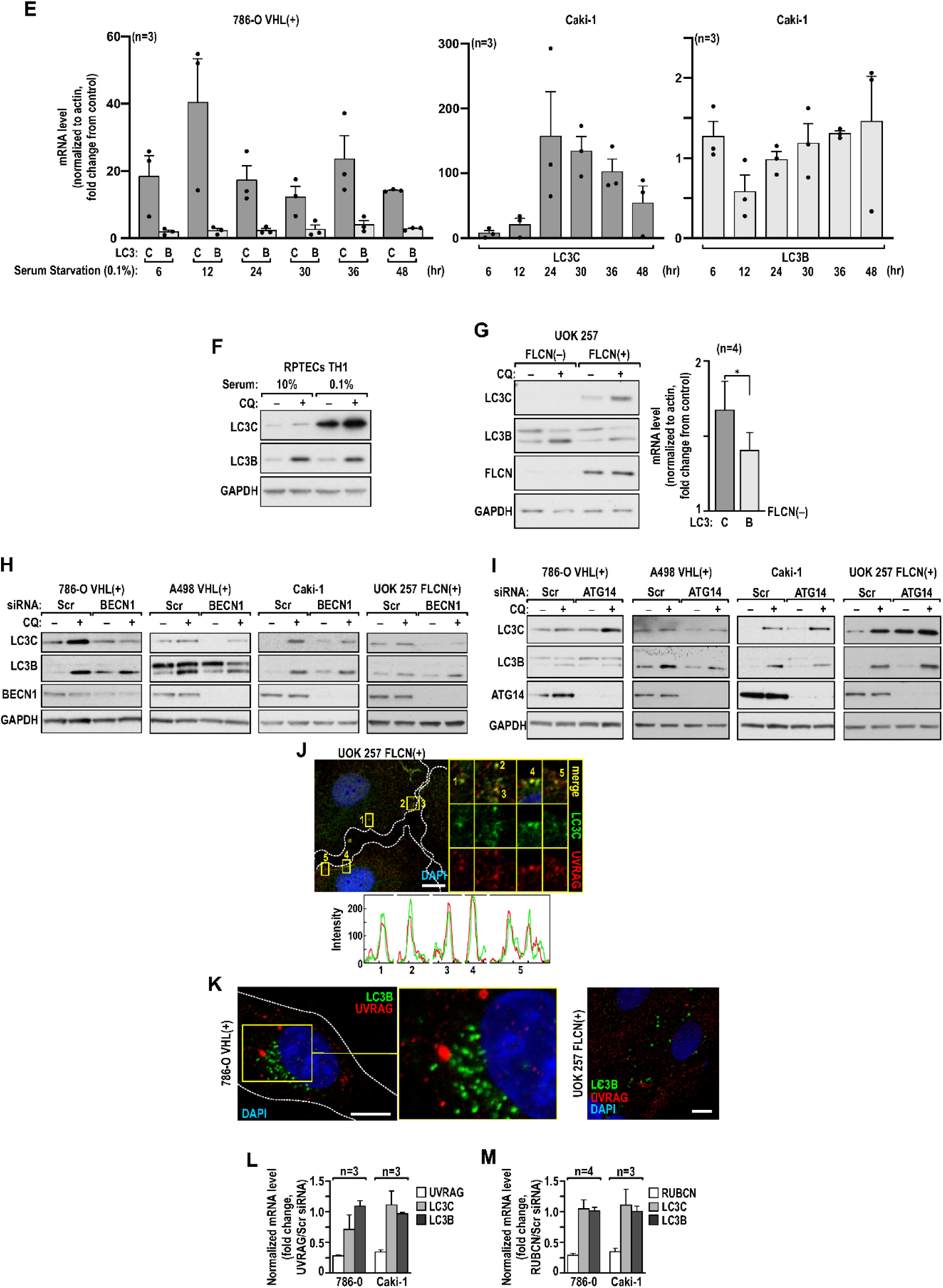
LC3C and LC3B are independent autophagic programs; LC3C is regulated by UVRAG and RUBCN. (A) Specificity of the anti-LC3C antibody in Caki-1 cells expressing endogenous LC3C or with stably knocked down LC3C. (B) Sequence of the LC3C C-terminal peptide is conserved among primates and humans. G126 undergoing lipidation is marked in yellow. (C) LC3C and LC3B similarly accumulate in response to 1 hr treatment with 100 µM chloroquine (CQ) or 200 nM Bafilomycin (BAFA1). (D) Immunofluorescence of LC3C and LC3B puncta with specific antibodies in the indicated cells treated without and with CQ for 1 hr to inhibit lysosomal processing. Inserts: quantification of puncta in the indicated number of cells. Higher magnification and quantification of the puncta shows that only a very small fraction colocalize. Mean ± SEM are shown. (E) Time course showing robust induction of LC3C but not LC3B mRNA in response to serum starvation (0.1% serum). For each time point data are normalized to mRNA measured in cells grown in 10% serum. Serum starvation induction of LC3C mRNA was significantly higher than induction of LC3B mRNA in 786-O VHL(+) cells at P=0.00138 and in Caki-1 at P=0.0106 (one way Anova). (F) LC3C, but not LC3B, is induced by serum starvation (48 hr, 0.1% serum) in human RPTEC cells. (G) Reexpression of FLCN in UOK 257 cells with lost *FLCN* induces LC3C autophagy. (H) LC3C flux is inhibited in response to BECN1 knockdown. (I) LC3C flux is not inhibited in response to ATG14 knockdown. (J) Colocalization of LC3C puncta with UVRAG. RGB profiles and split channels are shown. (K) Lack of colocalization of LC3B with UVRAG in the indicated cell lines. (L) RT-PCR shows that UVRAG knockdown does not affect LC3C or LC3B mRNA expression. (M) RT-PCR shows that RUBCN knockdown does not affect LC3C or LC3B mRNA expression. P values from two tailed unpaired t-test. Scale bars: 10 µm.

**Fig. S2.**
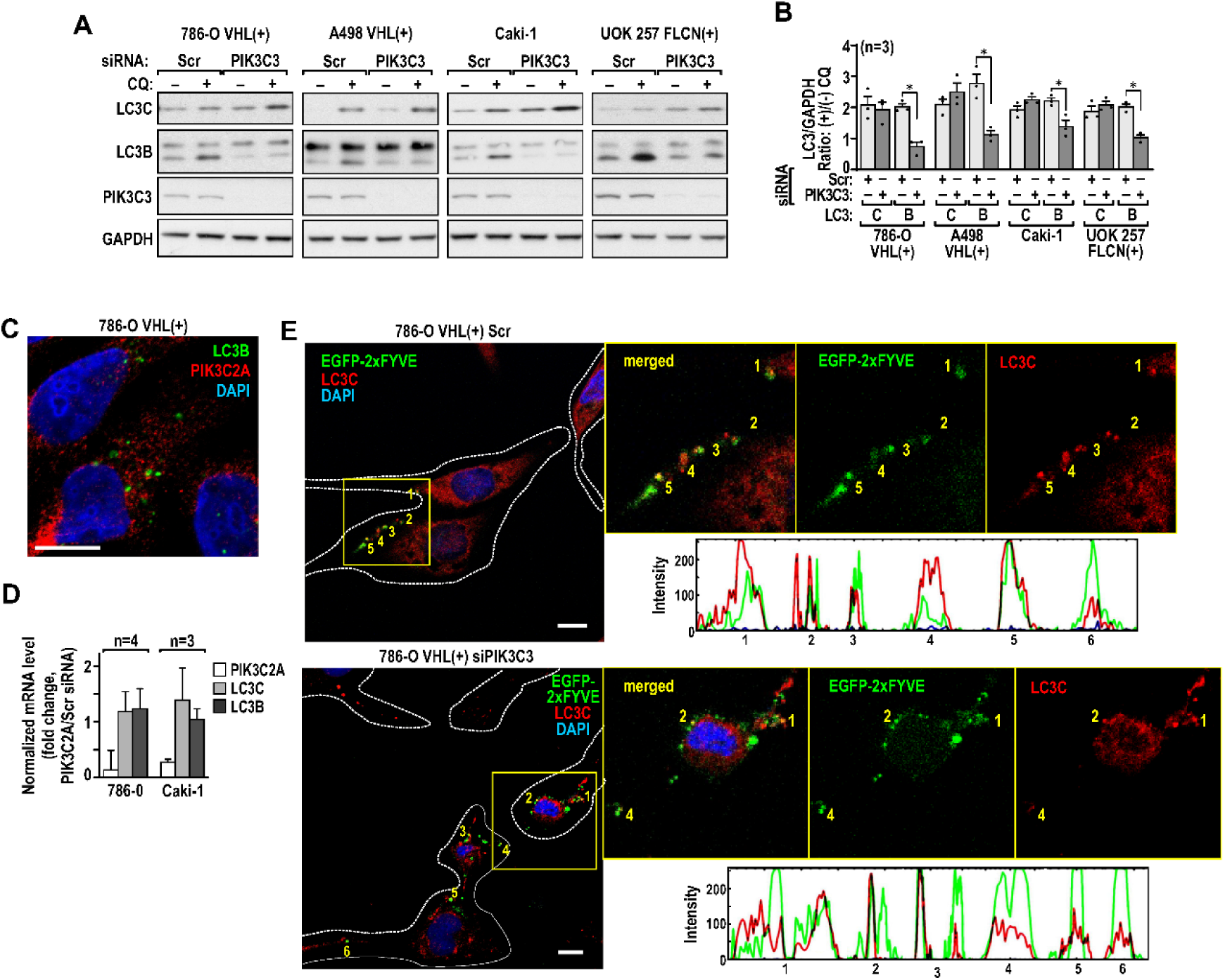
LC3C is regulated by PIK3C2A. (A) Immunoblots show that knockdown of PIK3C3 does not inhibit LC3C flux but is effective towards LC3B. (B) Quantification of LC3C and B accumulation presented as the ratio of their protein levels normalized to GAPDH in CQ(+) vs. CQ(-) in cells transfected with scramble or PIK3C3 siRNAs in the indicated cell lines. (C) Lack of colocalization of LC3B and PIK3C2A. (D) RT-PCR shows that PIK3C2A knockdown does not affect LC3C or LC3B mRNA expression. (E) Colocalization of LC3C with PI3P reporter, EGFP-2xFyve in cells with PIK3C3 knockdown. Means ± SEM are shown. Scale bars: 10 µm.

**Fig. S3.**
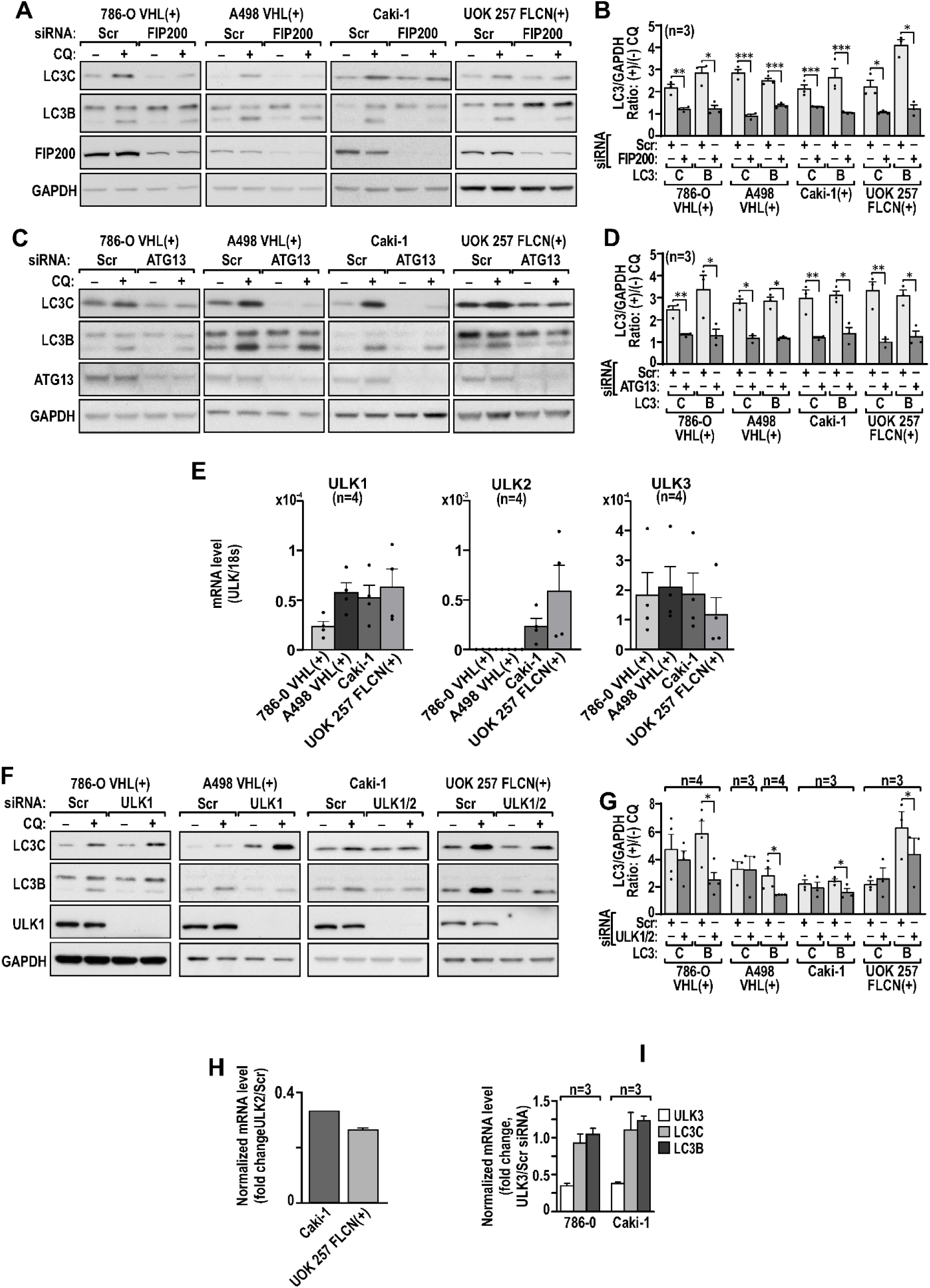
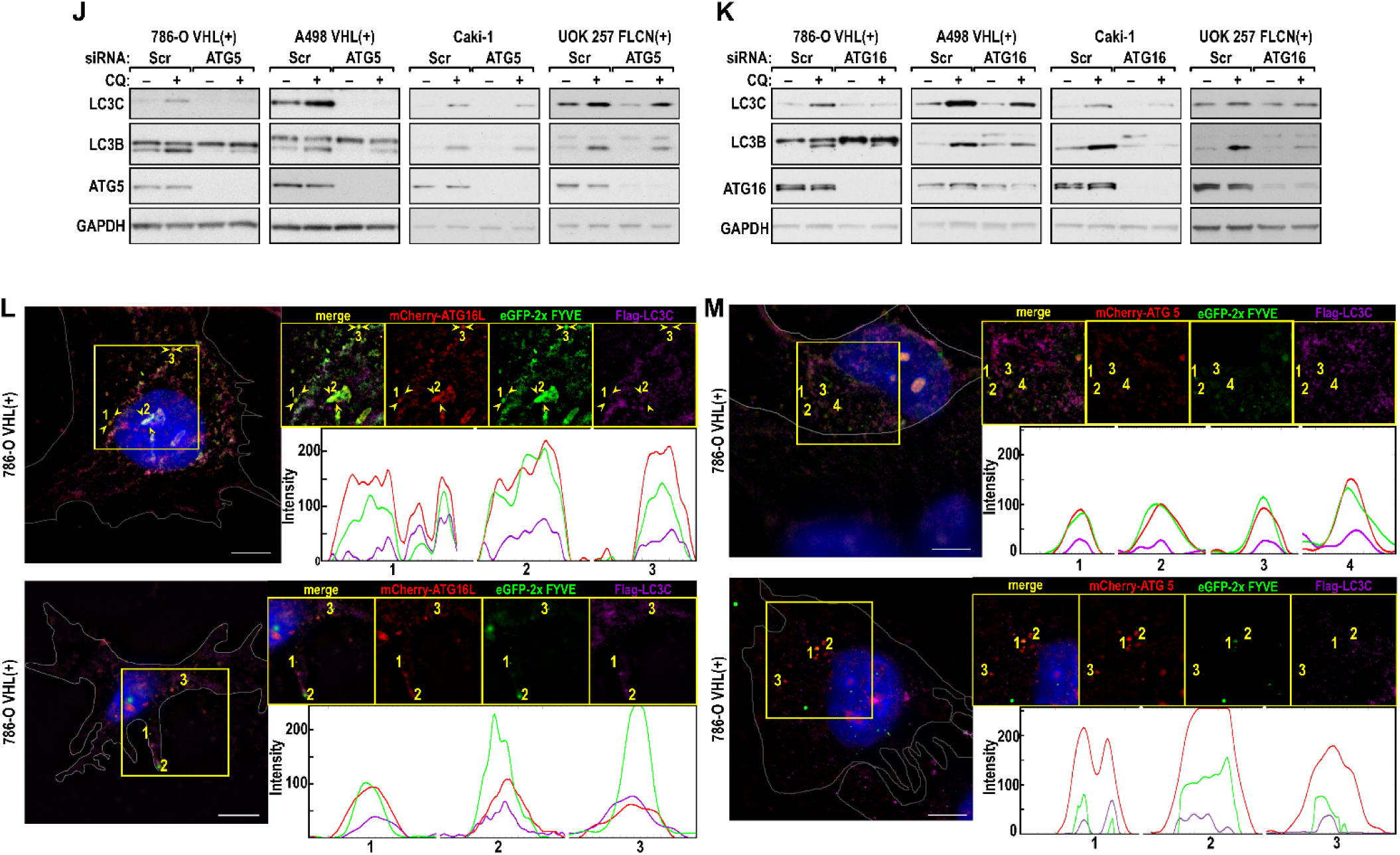
FIP200a and ATG13 are necessary for LC3Cactivity. (A) Immunoblots show a decrease in the CQ dependent accumulation of LC3C and LC3B in response to FIP200-KD in indicated cell lines. (B) Quantification of the LC3C and B accumulation presented as the ratio of their protein levels normalized to GAPDH in CQ(+) vs. CQ(-) in cells transfected with scramble or FIP200 siRNAs in the indicated cell lines. (C) Immunoblots show decrease in the CQ dependent accumulation of LC3C and LC3B in response to ATG13 KD in indicated cell lines. (D) Quantification of the LC3C and B accumulation presented as the ratio of their protein levels normalized to GAPDH in CQ(+) vs. CQ(-) in cells transfected with scramble or ATG13 siRNAs in the indicated cell lines. (E) RT PCR measurement of mRNA expression of ULK1, ULK2 and ULK3 in the indicated cell lines. (F) Immunoblots show effects of double knockdowns of ULK1 or ULK1/2 on LC3C and LC3B accumulation in response to CQ treatment. (G) Quantification of the LC3C and B accumulation presented as the ratio of their protein levels normalized to GAPDH in CQ(+) vs. CQ(-) in cells transfected with scramble or ULK1/2 siRNAs in the indicated cell lines. (H) ULK2 knockdown is measured by mRNA level. (I) Knockdown of ULK3 does not affect LC3C or LC3B mRNA. (J) Knockdown of ATG5 inhibits LC3C autophagy in the indicated cell lines. (K) Knockdown of ATG16 inhibits LC3C autophagy in the indicated cell lines. (L) Colocalization of LC3C with ATG16L and GFP-2xFyve puncta as analyzed by SIM. Examples of RGB profiles and split images are shown below. (M) Colocalization of LC3C with ATG and GFP-2xFyve puncta as analyzed by SIM. Examples of RGB profiles and split images are shown below. Means ± SEM are shown. P values are determined by unpaired two-tailed t-test. Scale bars: 10 µm.

**Fig. S4.**
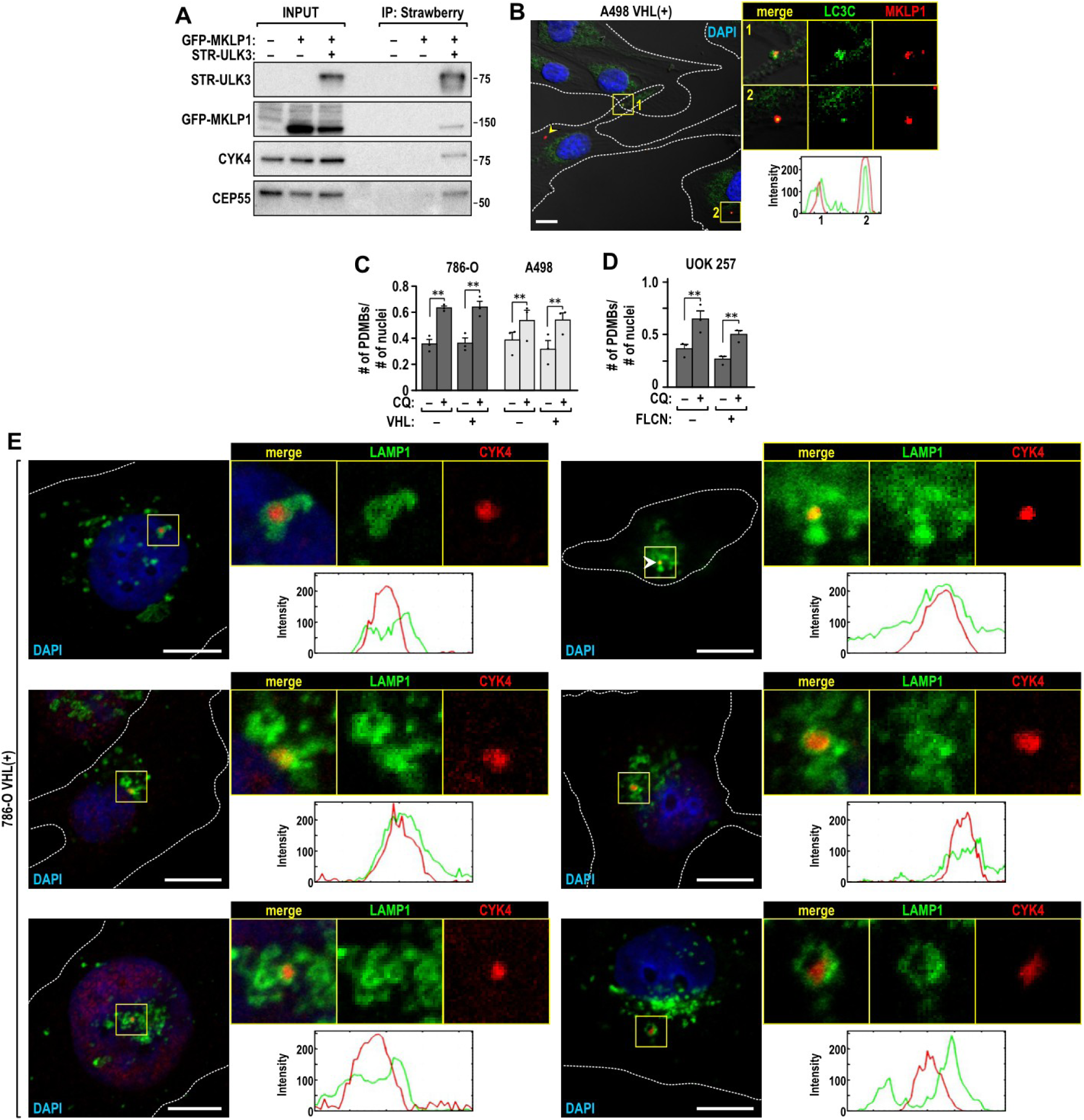
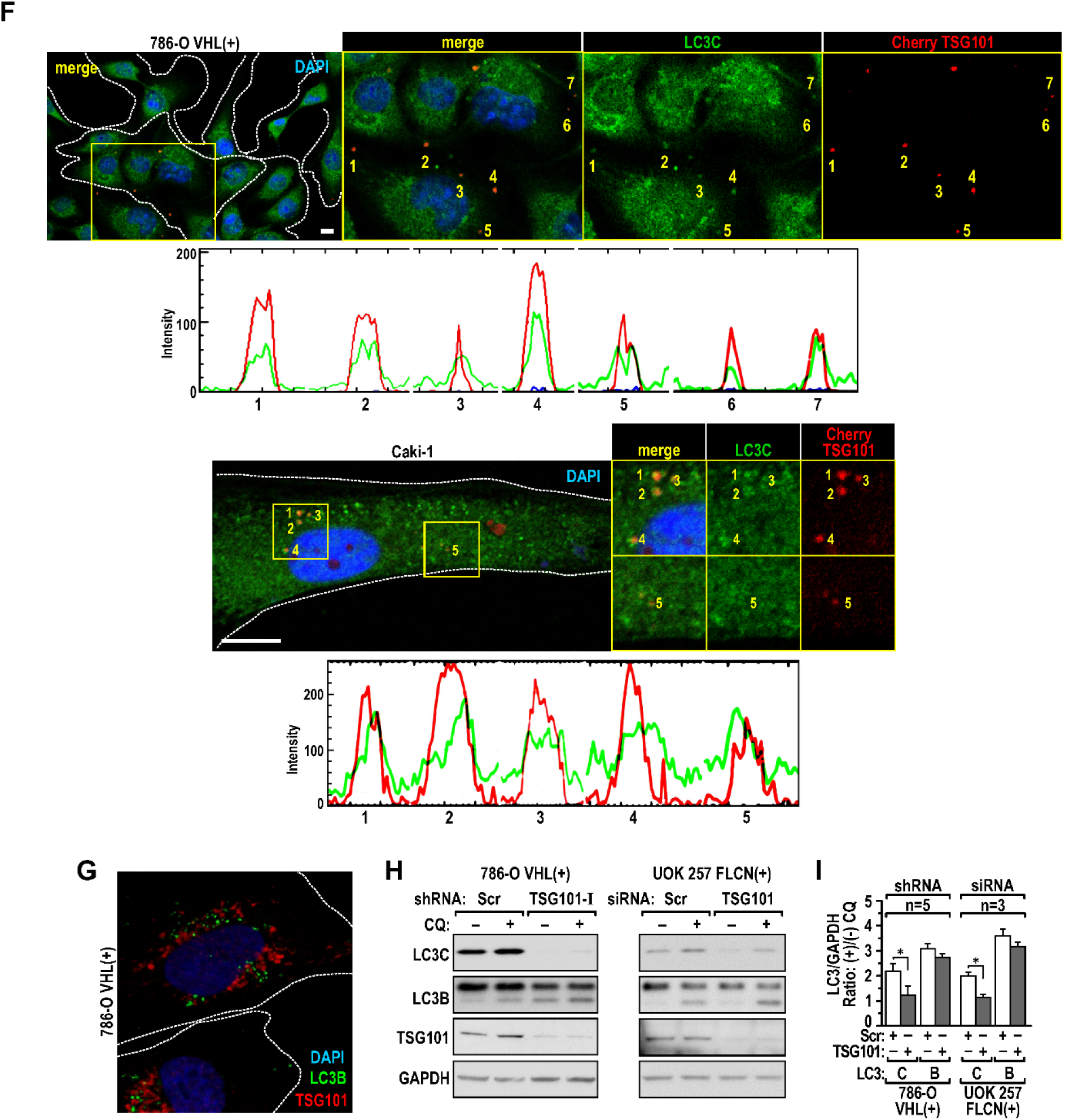
PDMBs are target of LC3C autophagy. (A) STR-ULK3 co-immunoprecipitates midbody proteins MKLP1, CYK4, and CEP55. (B) Colocalization of LC3C puncta with PDMBs in A498 VHL(+) cells. RGB profiles and split channels are shown. (C) VHL does not affect CQ-dependent degradation of PDMBs under 10% serum conditions. (D) FLCN does not affect CQ-dependent degradation of PDMBs under 10% serum conditions. In C and D quantification was performed as described in Fig. 4D. (E) Multiple examples of colocalization of PDMBs with LAMP1, a lysosomal marker. RGB profiles and split channels are shown. (F) Cherry-TSG101 colocalizes with endogenous LC3C in 786-O VHL(+) and Caki-1 cells. RGB profiles and split channels are shown. (G) Cherry-TSG101 does not colocalize with endogenous LC3B. (H) Knockdown of TSG101 inhibits LC3C autophagy. (I) Quantification of the LC3C and B accumulation presented as the ratio of their protein levels normalized to GAPDH in CQ(+) vs. CQ(-) in cells transfected with scr or TSG101 si or shRNA in the indicated cell lines. P value calculated by unpaired, two-tailed t-test. Scale bars:10 µm

**Fig. S5.**
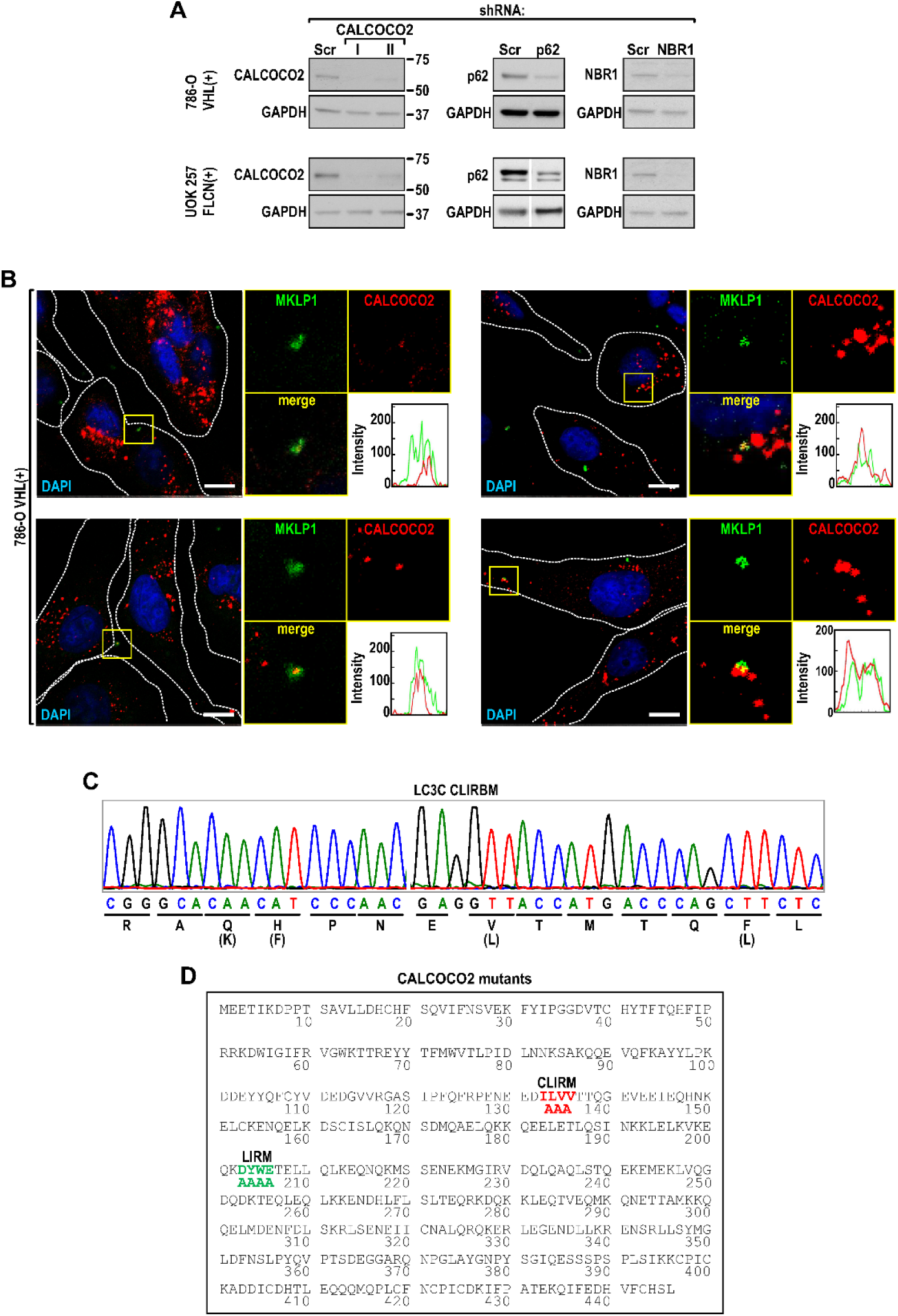
Selectivity of LC3C autophagy requires LIR binding motif and LIR on CALCOCO2. (A) Immunoblots showing knockdowns of respective cargo receptors. (B) Representative examples of colocalization of PDMBs with endogenous CALCOCO2. RGB profiles and split channels are shown. MKLP1 is used as a PDMB marker. (C) Sequence of LC3C with mutations in CLIR binding site (CLRBM). The wild type sequence indicated in parenthesis. (D) Mutations of CLIR and LIR motifs in CALCOCO2. Optimized CALCOCO2 was a synthetic gene. Scale bars: 5 µm.

**Fig. S6.**
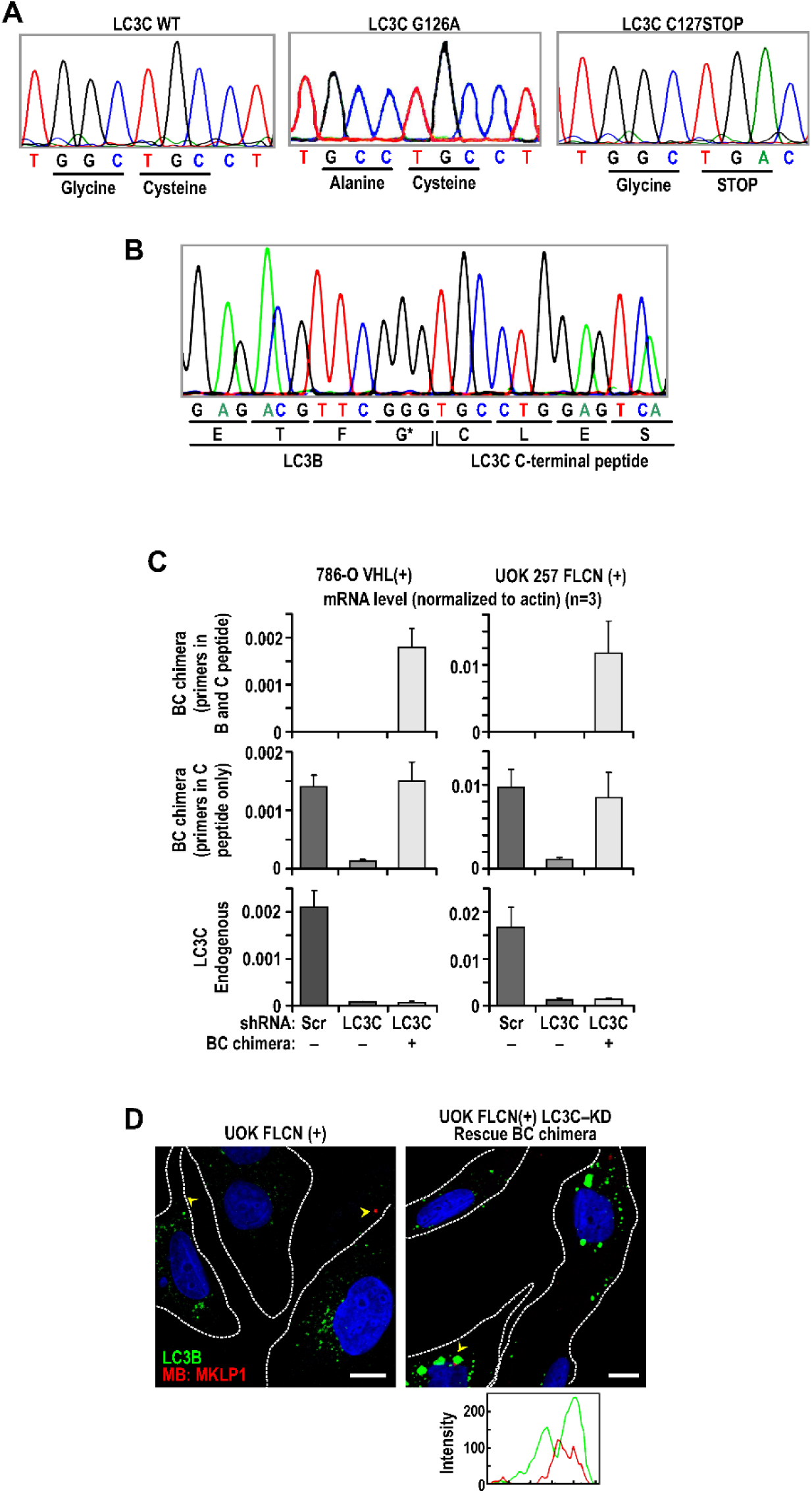
Role of LC3C peptide in regulation of LC3C autophagy. (A) Sequence of the G126A, C127 Stop mutants (B) Sequence of BC chimera (C) RT PCR quantification of mRNA expression levels using primers specific for the BC chimeras only (top), primers detecting levels of mRNAs containing C-terminal peptide only (middle) and primers specific for LC3C only (bottom). Experiments were done in two indicated cell lines (D) Representative examples of immunofluorescence experiments showing lack of colocalization of PDMBs with LC3B in cells with LC3C knockdown (left), but appearance of close proximity and colocalization in cells expressing BC chimeras (right) in UOK 257 cells. RGB profiles are shown.

**Table S1.**
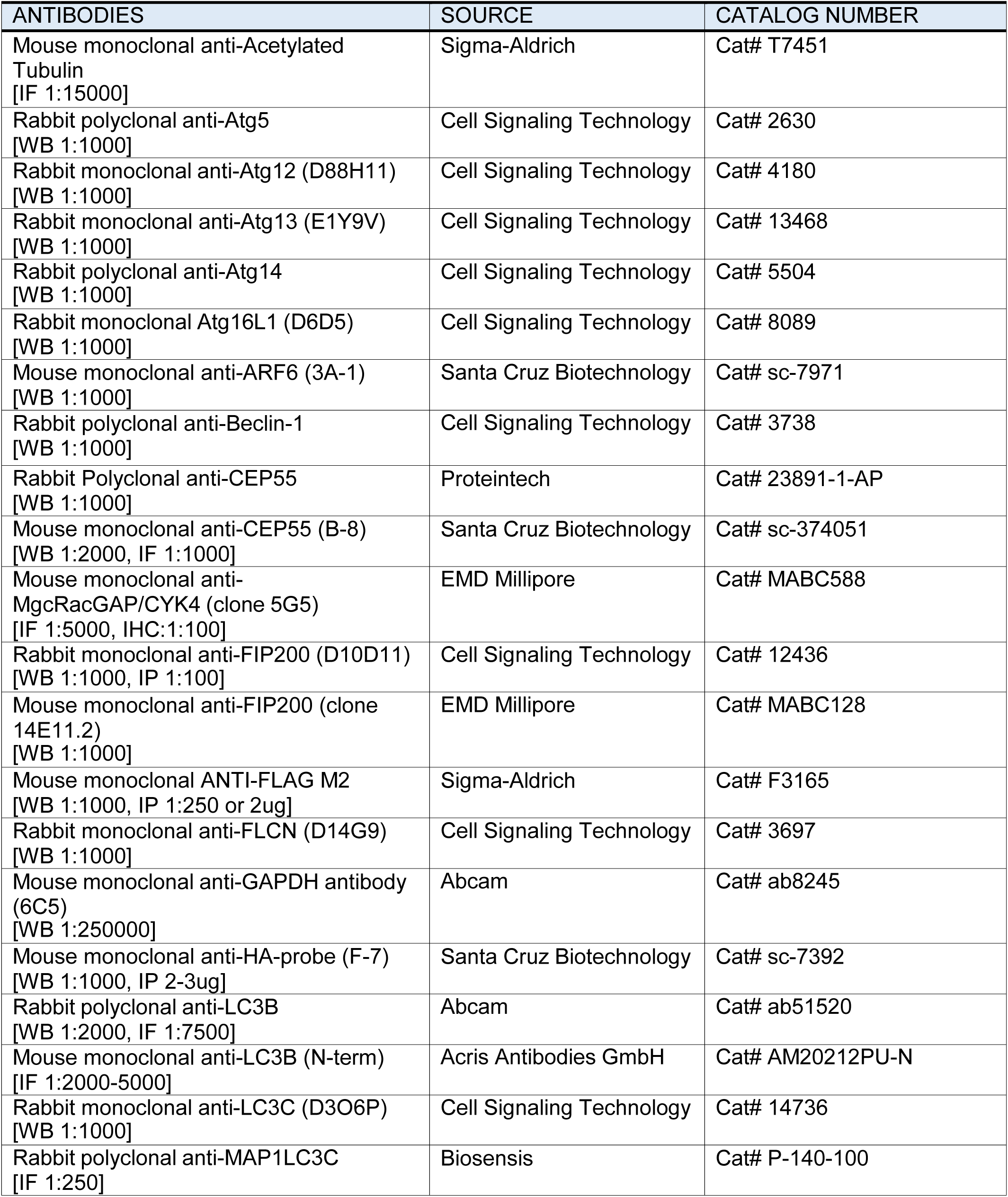
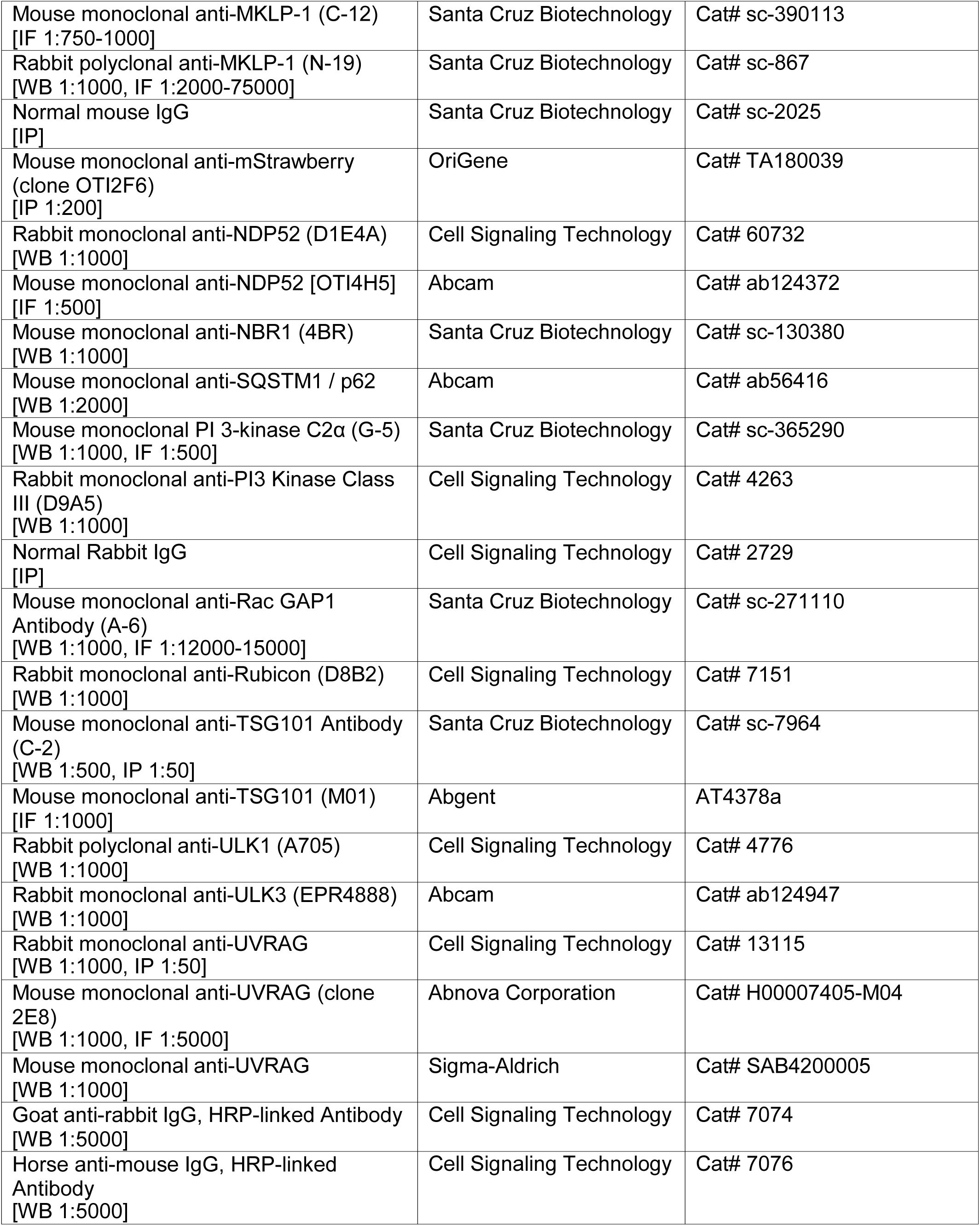
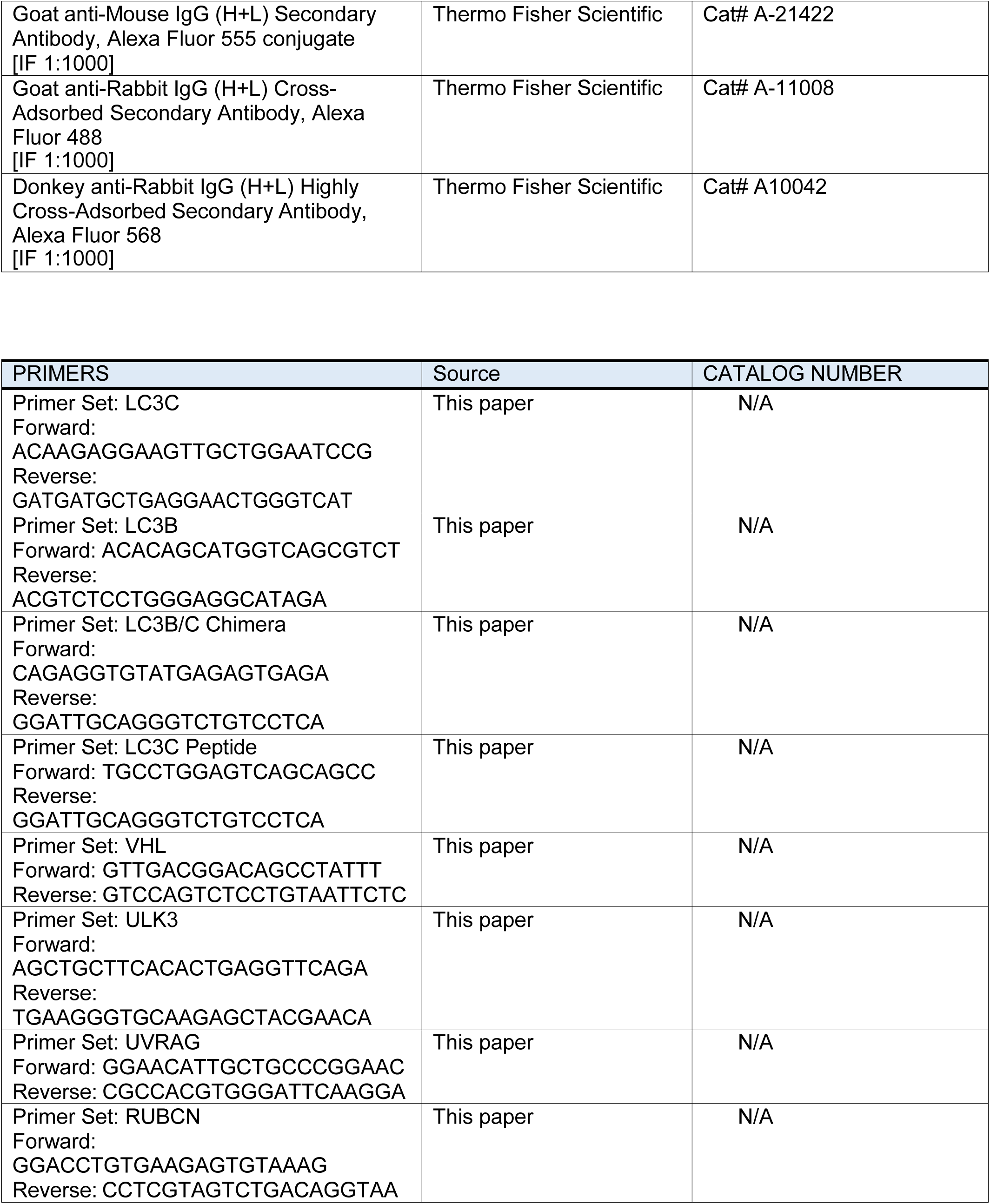
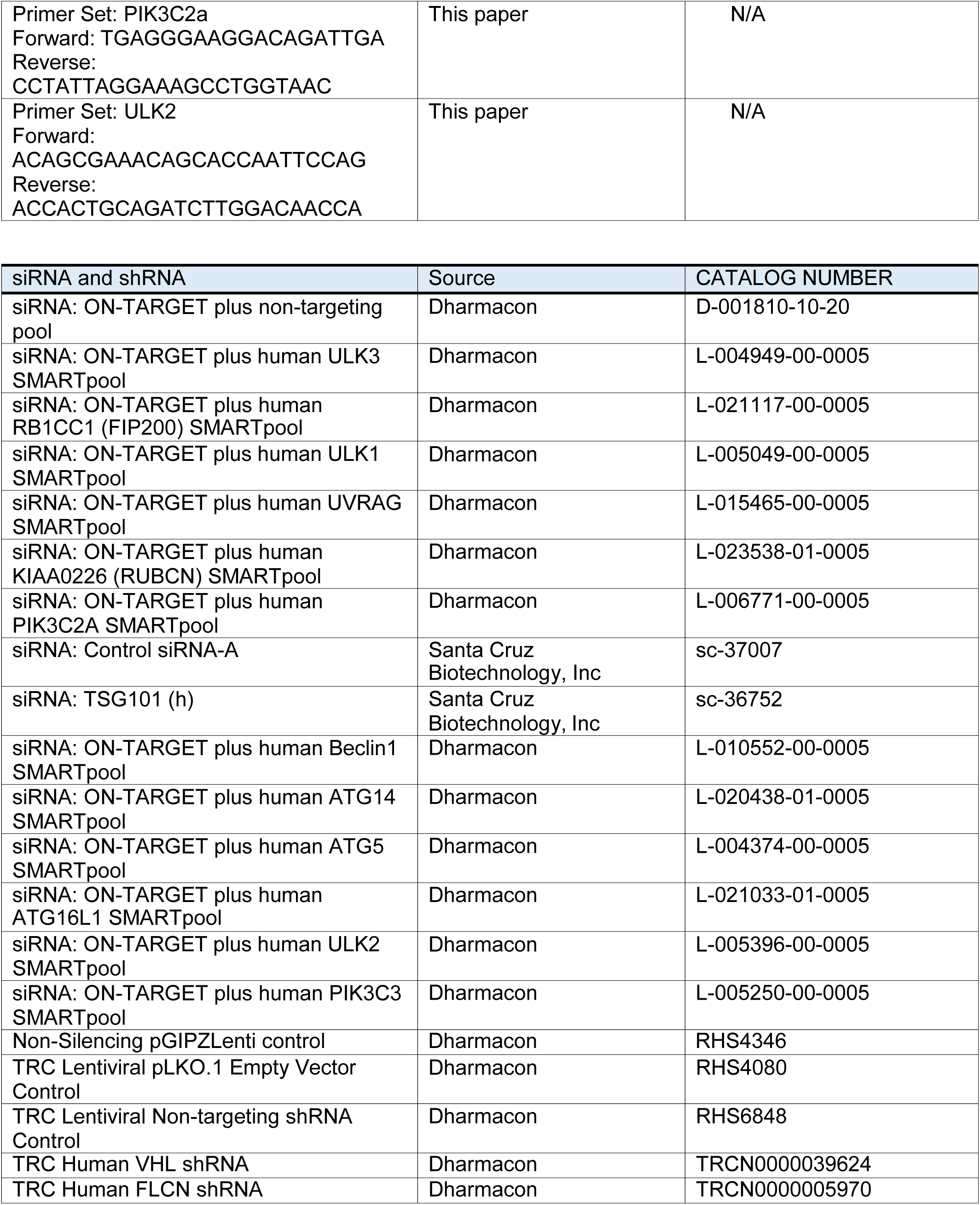
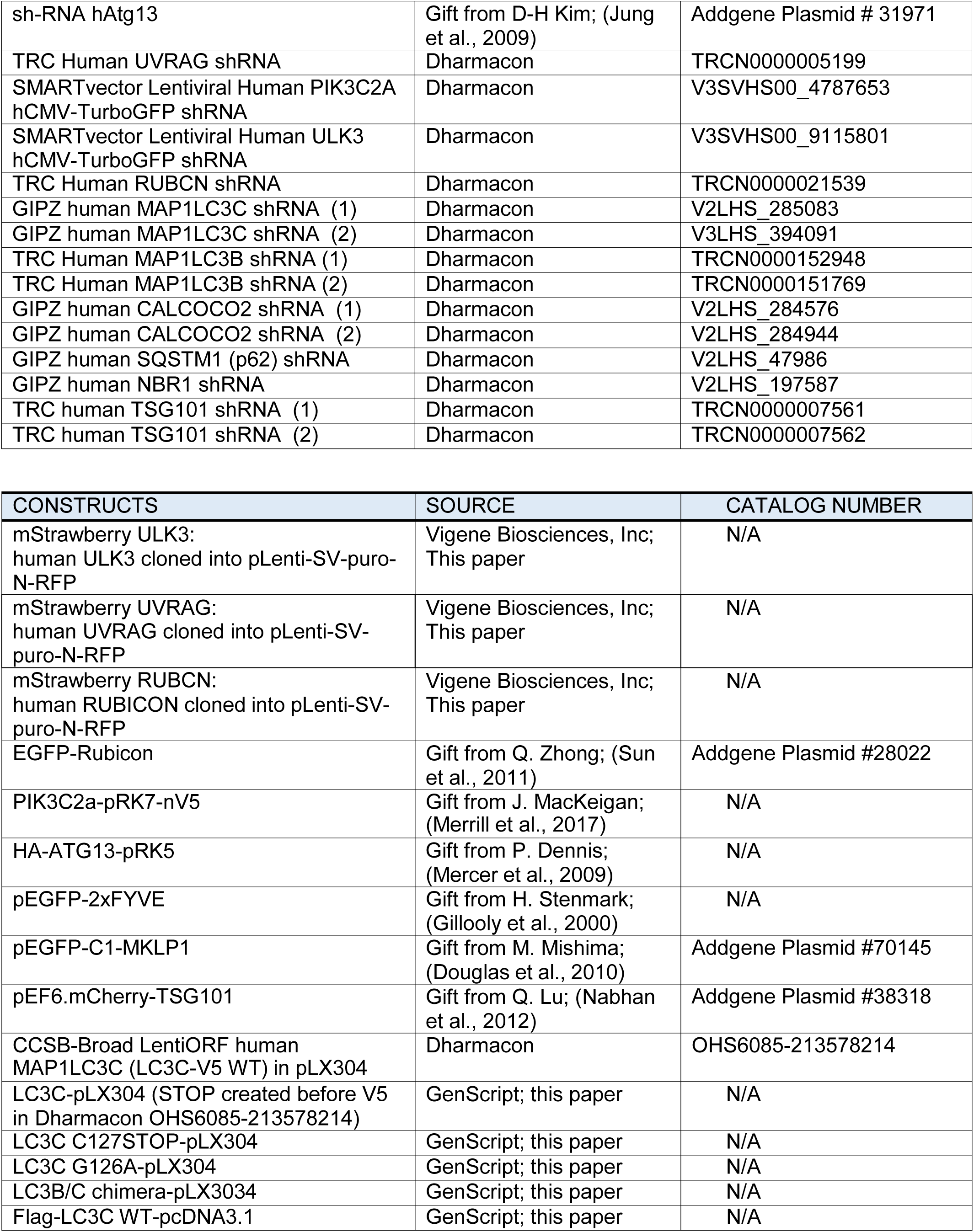
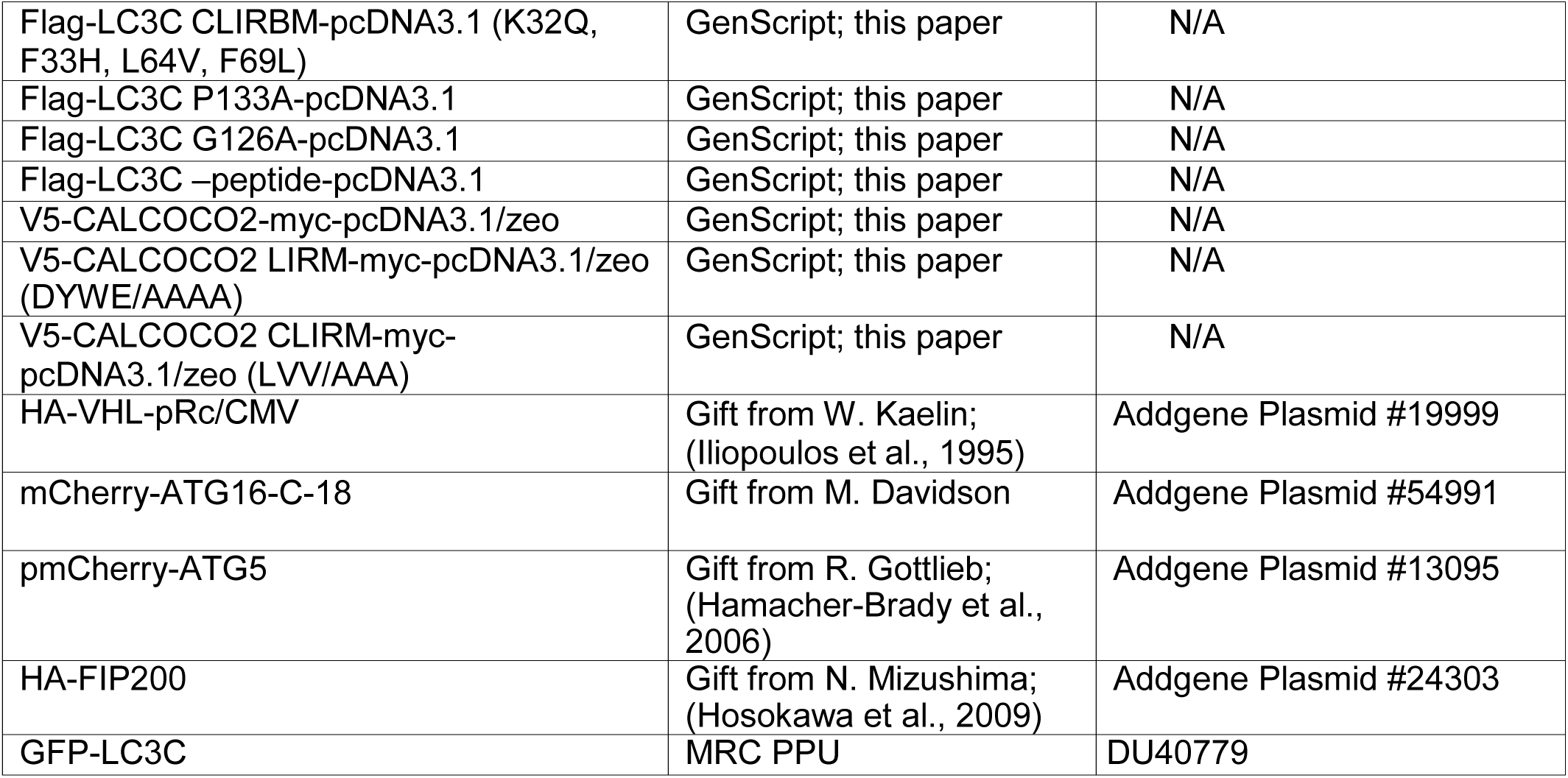
List of reagents used in the study.

## REFERENCES

Bastola, P., Stratton, Y., Kellner, E., Mikhaylova, O., Yi, Y., Sartor, M.A., Medvedovic, M., Biesiada, J., Meller, J., and Czyzyk-Krzeska, M.F. (2013). Folliculin contributes to VHL tumor suppressing activity in renal cancer through regulation of autophagy. PloS one 8, e70030.

Bell, E.S., Coelho, P.P., Ratcliffe, C.D.H., Rajadurai, C.V., Peschard, P., Vaillancourt, R., Zuo, D., and Park, M. (2019). LC3C-Mediated Autophagy Selectively Regulates the Met RTK and HGF-Stimulated Migration and Invasion. Cell reports 29, 4053–4068.e4056.

Caballe, A., Wenzel, D.M., Agromayor, M., Alam, S.L., Skalicky, J.J., Kloc, M., Carlton, J.G., Labrador, L., Sundquist, W.I., and Martin-Serrano, J. (2015). ULK3 regulates cytokinetic abscission by phosphorylating ESCRT-III proteins. eLife 4, e06547.

Crowell, E.F., Gaffuri, A.L., Gayraud-Morel, B., Tajbakhsh, S., and Echard, A. (2014). Engulfment of the midbody remnant after cytokinesis in mammalian cells. Journal of cell science 127, 3840–3851.

Crowell, E.F., Tinevez, J.Y., and Echard, A. (2013). A simple model for the fate of the cytokinesis midbody remnant: implications for remnant degradation by autophagy. BioEssays : news and reviews in molecular, cellular and developmental biology 35, 472–481.

Fazeli, G., Trinkwalder, M., Irmisch, L., and Wehman, A.M. (2016). C. elegans midbodies are released, phagocytosed and undergo LC3-dependent degradation independent of macroautophagy. Journal of cell science 129, 3721–3731.

Fazeli, G., and Wehman, A.M. (2017). Safely removing cell debris with LC3-associated phagocytosis. Biology of the cell 109, 355–363.

Galluzzi, L., Baehrecke, E.H., Ballabio, A., Boya, P., Bravo-San Pedro, J.M., Cecconi, F., Choi, A.M., Chu, C.T., Codogno, P., Colombo, M.I., et al. (2017). Molecular definitions of autophagy and related processes. The EMBO journal 36, 1811–1836.

Hall, D.P., Cost, N.G., Hegde, S., Kellner, E., Mikhaylova, O., Stratton, Y., Ehmer, B., Abplanalp, W.A., Pandey, R., Biesiada, J., et al. (2014). TRPM3 and miR-204 establish a regulatory circuit that controls oncogenic autophagy in clear cell renal cell carcinoma. Cancer cell 26, 738–753.

Herhaus, L., Bhaskara, R.M., Lystad, A.H., Gestal-Mato, U., Covarrubias-Pinto, A., Bonn, F., Simonsen, A., Hummer, G., and Dikic, I. (2020). TBK1-mediated phosphorylation of LC3C and GABARAP-L2 controls autophagosome shedding by ATG4 protease. EMBO Rep 21, e48317.

Kuo, T.C., Chen, C.T., Baron, D., Onder, T.T., Loewer, S., Almeida, S., Weismann, C.M., Xu, P., Houghton, J.M., Gao, F.B., et al. (2011). Midbody accumulation through evasion of autophagy contributes to cellular reprogramming and tumorigenicity. Nature cell biology 13, 1214–1223.

Lai, S.C., and Devenish, R.J. (2012). LC3-Associated Phagocytosis (LAP): Connections with Host Autophagy. Cells 1, 396–408.

Le Guerroue, F., Eck, F., Jung, J., Starzetz, T., Mittelbronn, M., Kaulich, M., and Behrends, C. (2017). Autophagosomal Content Profiling Reveals an LC3C-Dependent Piecemeal Mitophagy Pathway. Molecular cell 68, 786–796.e786.

Liang, C., Feng, P., Ku, B., Dotan, I., Canaani, D., Oh, B.H., and Jung, J.U. (2006). Autophagic and tumour suppressor activity of a novel Beclin1-binding protein UVRAG. Nature cell biology 8, 688–699.

Liang, C., Lee, J.S., Inn, K.S., Gack, M.U., Li, Q., Roberts, E.A., Vergne, I., Deretic, V., Feng, P., Akazawa, C., et al. (2008). Beclin1-binding UVRAG targets the class C Vps complex to coordinate autophagosome maturation and endocytic trafficking. Nature cell biology 10, 776–787.

Madjo, U., Leymarie, O., Fremont, S., Kuster, A., Nehlich, M., Gallois-Montbrun, S., Janvier, K., and Berlioz-Torrent, C. (2016). LC3C Contributes to Vpu-Mediated Antagonism of BST2/Tetherin Restriction on HIV-1 Release through a Non-canonical Autophagy Pathway. Cell reports 17, 2221–2233.

Martinez, J., Cunha, L.D., Park, S., Yang, M., Lu, Q., Orchard, R., Li, Q.Z., Yan, M., Janke, L., Guy, C., et al. (2016). Noncanonical autophagy inhibits the autoinflammatory, lupus-like response to dying cells. Nature 533, 115–119.

Martinez, J., Malireddi, R.K., Lu, Q., Cunha, L.D., Pelletier, S., Gingras, S., Orchard, R., Guan, J.L., Tan, H., Peng, J., et al. (2015). Molecular characterization of LC3-associated phagocytosis reveals distinct roles for Rubicon, NOX2 and autophagy proteins. Nature cell biology 17, 893–906.

Matsunaga, K., Saitoh, T., Tabata, K., Omori, H., Satoh, T., Kurotori, N., Maejima, I., Shirahama-Noda, K., Ichimura, T., Isobe, T., et al. (2009). Two Beclin 1-binding proteins, Atg14L and Rubicon, reciprocally regulate autophagy at different stages. Nature cell biology 11, 385–396.

Merrill, N.M., Schipper, J.L., Karnes, J.B., Kauffman, A.L., Martin, K.R., and MacKeigan, J.P. (2017). PI3K-C2alpha knockdown decreases autophagy and maturation of endocytic vesicles. PloS one 12, e0184909.

Mikhaylova, O., Stratton, Y., Hall, D., Kellner, E., Ehmer, B., Drew, A.F., Gallo, C.A., Plas, D.R., Biesiada, J., Meller, J., et al. (2012). VHL-regulated MiR-204 suppresses tumor growth through inhibition of LC3B-mediated autophagy in renal clear cell carcinoma. Cancer cell 21, 532–546.

Pohl, C., and Jentsch, S. (2009). Midbody ring disposal by autophagy is a post-abscission event of cytokinesis. Nature cell biology 11, 65–70.

Ravenhill, B.J., Boyle, K.B., von Muhlinen, N., Ellison, C.J., Masson, G.R., Otten, E.G., Foeglein, A., Williams, R., and Randow, F. (2019). The Cargo Receptor NDP52 Initiates Selective Autophagy by Recruiting the ULK Complex to Cytosol-Invading Bacteria. Molecular cell 74, 320–329.e326.

Salzmann, V., Chen, C., Chiang, C.Y., Tiyaboonchai, A., Mayer, M., and Yamashita, Y.M. (2014). Centrosome-dependent asymmetric inheritance of the midbody ring in Drosophila germline stem cell division. Molecular biology of the cell 25, 267–275.

Sanjuan, M.A., Dillon, C.P., Tait, S.W., Moshiach, S., Dorsey, F., Connell, S., Komatsu, M., Tanaka, K., Cleveland, J.L., Withoff, S., et al. (2007). Toll-like receptor signalling in macrophages links the autophagy pathway to phagocytosis. Nature 450, 1253–1257.

Stadel, D., Millarte, V., Tillmann, K.D., Huber, J., Tamin-Yecheskel, B.C., Akutsu, M., Demishtein, A., Ben-Zeev, B., Anikster, Y., Perez, F., et al. (2015). TECPR2 Cooperates with LC3C to Regulate COPII-Dependent ER Export. Molecular cell 60, 89–104.

Thoresen, S.B., Pedersen, N.M., Liestol, K., and Stenmark, H. (2010). A phosphatidylinositol 3-kinase class III sub-complex containing VPS15, VPS34, Beclin 1, UVRAG and BIF-1 regulates cytokinesis and degradative endocytic traffic. Experimental cell research 316, 3368–3378.

von Muhlinen, N., Akutsu, M., Ravenhill, B.J., Foeglein, A., Bloor, S., Rutherford, T.J., Freund, S.M., Komander, D., and Randow, F. (2012). LC3C, bound selectively by a noncanonical LIR motif in NDP52, is required for antibacterial autophagy. Molecular cell 48, 329–342.

White, E. (2015). The role for autophagy in cancer. The Journal of clinical investigation 125, 42–46.

White, E.A., and Glotzer, M. (2012). Centralspindlin: at the heart of cytokinesis. Cytoskeleton (Hoboken, NJ) 69, 882–892.

Zaffagnini, G., and Martens, S. (2016). Mechanisms of Selective Autophagy. Journal of molecular biology 428, 1714–1724.

